# The Myo2 adaptor Ldm1 and its receptor Ldo16 mediate actin-dependent lipid droplet motility

**DOI:** 10.1101/2025.02.21.639583

**Authors:** Xue-Tong Zhao, Duy Trong Vien Diep, Louis Percifull, Rebecca Martina Fausten, Marie Hugenroth, Pascal Höhne, Bianca Marie Esch, Javier Collado, Jenny Keller, Stephan Wilmes, Mike Wälte, Daniel Kümmel, Christian Schuberth, Rubén Fernández-Busnadiego, Florian Fröhlich, Roland Wedlich-Söldner, Maria Bohnert

**Author notes:** These authors contributed equally to this work (Duy Trong Vien Diep, Louis Percifull, Rebecca Martina Fausten, Marie Hugenroth, Pascal Höhne). Correspondence should be addressed to (M.B.).

## Abstract

Organelle motility enables strategic cellular reorganizations. In yeast, this process depends on the actin cytoskeleton, type V myosin motor proteins, and organelle-specific myosin adaptor proteins. While the myosin adaptors for most organelles are known, the coupling of myosin to lipid droplets (LDs), the cellular lipid storage organelles, remained enigmatic. Using genome-wide screening, we identified Ldm1 (Lipid Droplet Motility 1/Yer085c) as a myosin adaptor. Ldm1 binds to the globular tail domain of the myosin Myo2 and to the LD surface protein Ldo16 to enable actin-dependent LD motility. Ldo16 has additional roles in LD contact sites to the vacuole and the ER, suggesting a coordination of LD motility and organelle tethering. Ldm1 has a second role in mitochondrial transport and elevated Ldm1 levels rescue defects of the mitochondrial Myo2-adaptors Mmr1/Ypt11. Our work identifies the molecular machinery for LD motility and contributes to a comprehensive understanding of acto-myosin-based cellular reorganization.

## INTRODUCTION

The interior of eukaryotic cells contains a set of organelles that are spatially highly organized. Organelle motility is crucial for strategic reorganizations of cellular content in response to metabolic cues and environmental stimuli ^1^, during differentiation ^2^ and in cell division ^3^. In *Saccharomyces cerevisiae* (from hereon: yeast), organelle motility depends on the actin cytoskeleton ^4^ with the help of the class V myosin motors Myo2 ^5–10^ and Myo4 ^11^. Specificity is achieved by a set of myosin adaptor proteins, which mediate myosin binding to the surfaces of the different organelles ^12^. The Myo2 adaptor protein for peroxisomes is Inp2 ^13^, for the vacuole (lysosome-like organelle) Vac17 ^14,15^, and for mitochondria two adaptors are known, Mmr1 and Ypt11 ^16–18^. Ypt11 is multifunctional and additionally promotes transport of the late Golgi ^19^. Myo4 mediates motility of the endoplasmic reticulum (ER), together with its adaptor She3 ^11^. Some myosin adaptors bind directly to the respective organelle membrane, such as the peroxisome adaptor Inp2 ^13^, while others require an organelle surface receptor for organelle attachment, such as Vac17 that forms a complex with the vacuole surface receptor Vac8 ^14,15^.

A further motile organelle is the lipid droplet (LD), the cellular lipid storage organelle, but the molecular basis for LD motility is currently incompletely understood ^20,21^. By storing variable amounts of lipids, LDs ensure survival during nutrient shortage and provide a safe space for toxic lipid species. LD dysfunctions are linked to human diseases including obesity, diabetes, fatty liver disease, infectious diseases and lipodystrophy ^22,23^. LDs are formed at the membrane of the ER, where enzymes for the synthesis of neutral storage lipids such as triglycerides and sterol esters are located ^24,25^. These neutral lipids form oil lenses between the leaflets of the ER phospholipid bilayer, which grow and ultimately bud toward the cytoplasm ^26^. Therefore, LDs have a special architecture, consisting of a central neutral lipid core and an outer phospholipid monolayer. The phospholipid monolayer houses the LD surface proteome that comprises lipid metabolism enzymes, proteins involved in LD formation, turnover, morphology, and communication with other organelles, and proteins of unknown function ^27,28^. Due to their key roles in lipid storage and metabolism, LDs are tightly connected to all other organelles via membrane contact sites for collaborative functions in lipid handling ^29–31^. In mammalian cells, both microtubule- and actin-based cytoskeletal structures are involved in remodeling of the LD contact site landscape and adaptation of LD metabolism ^32–36^. Across species, it is well established that LDs are motile in a manner dependent on the cytoskeleton, but comparatively little is known about LD-specific motor protein adaptors ^20,21^. In yeast, it has been shown that LDs move toward developing daughter cells during cell division ^37^ dependent on Myo2 ^10^, but it has been unclear how this motor protein is attached to the LD surface to drive organelle motility.

Here, we used systematic genetic and proteomic approaches to uncover the unknown LD-specific motility factors in yeast. We identified the protein of unknown function Yer085c (now Lipid Droplet Motility 1, Ldm1) as the missing Myo2 adaptor for LDs. Ldm1 interacts with the globular tail domain of Myo2 and binds to LDs via the LD surface receptor Ldo16, a multifunctional LD protein that is also involved in seipin-mediated LD biogenesis ^38–40^ and in formation of vacuole-LD contact sites ^41,42^. We find that Ldm1 is a bifunctional Myo2 adaptor that additionally affects motility of mitochondria. Ldm1 overexpression restores viability of mutants of the mitochondrial Myo2 adaptors Mmr1 and Ypt11. Ldm1-dependent motility of LDs and mitochondria can be genetically uncoupled by deletion of *LDO16*. Our findings answer the long-standing question how LD motility is achieved on a molecular level and pave the way to a comprehensive understanding of acto-myosin based cellular reorganization.

## RESULTS

### A genome-wide screen identifies determinants of cellular LD distribution and abundance

LDs are motile organelles, but the molecular basis for LD motility is incompletely understood ^10^. We grew cells expressing the fluorescent LD surface protein Erg6-mCherry to logarithmic growth phase in synthetic complete media and used time-resolved microscopy to assess LD motility. We observed different motility states for LDs (Figure 1A). At any given timepoint, a part of the LD pool was largely immotile, visible as small, circular foci in maximal projections, and as straight vertical lines in kymographs (Figure 1A, left maximal projection and kymograph 1). Periodically, LDs switched to a state of directional motility, visible as elongated streaks in maximal projections, and as horizontal line shifts in kymographs (Figure 1A, left maximal projection, kymographs 2 and 3, and bottom panel). Treatment of cells with latrunculin B to inhibit polymerization of actin filaments blocked directional LD motility (Figure 1A, right), consistent with previous reports that identified the type V myosin Myo2 as a player involved in LD motility ^10^.

**Figure 1.**
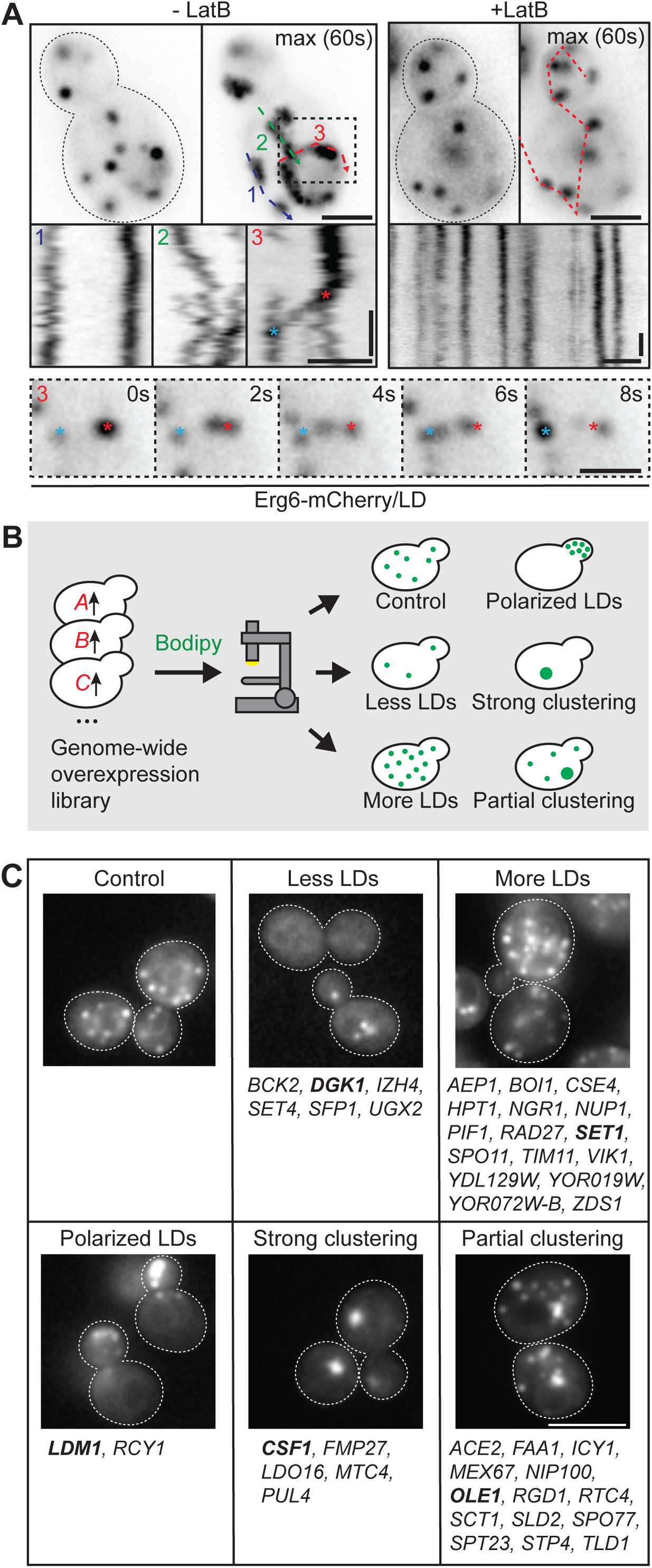
A microscopy-based screen identifies effectors of LD distribution and abundance. (A) Cells expressing the LD marker Erg6-mCherry were treated for 2 hours with 200 µM of Latrunculin B (right, +LatB) or left untreated (left, -LatB), and analyzed by time-resolved microscopy at 2 frames per second (s). Single frames, maximal projections of 60 s, and kymographs along color coded dotted lines are shown. Bottom row shows an LD motility event (movement from red to blue asterisk) from the -LatB cell (dashed outline). Scale bars, 2 µm. Time bars, 10 s. (B) Schematic representation of a screen for genes affecting LD distribution and abundance. A genome-wide collection of strains carrying alleles for overexpression of mCherry-tagged proteins from a strong *TEF2* promoter was analyzed by staining with the neutral lipid dye BODIPY493/503 and automated microscopy. Five phenotypical categories of abnormal LD distribution and abundance were observed: less LDs, more LDs, strong LD clustering, partial LD clustering, and polarized LD localization. (C) Lists of genes that, when overexpressed, induced LD distribution and abundance phenotypes described in (B). Genes shown in representative images are labeled in bold. Scale bar, 5 µm. Also see Table S1.

Yeast cells undergo asymmetric cell divisions, in which daughter cells, also termed buds, grow from one end of the mother cells. In this process, actin cables assemble in a polarized manner to allow for type V myosin-dependent vectorial delivery of secretory vesicles and organelles from the mother cell to the bud ^43^. As LD motility depends on actin (Figure 1A) and Myo2 ^10^, we reasoned that in dividing cells, overexpression of proteins involved in LD motility should result in an enhanced bud-directed LD transport and consequently an accumulation of LDs in buds. We therefore designed an overexpression-based unbiased screen using a genome-wide mutant collection, in which every protein was expressed under the control of the strong, constitutive *TEF2* promoter and additionally fused to an N-terminal mCherry moiety ^44,45^. This collection of ∼6000 strains was cultured to logarithmic growth phase, followed by staining with the neutral lipid dye BODIPY493/503 and analysis by automated microscopy (Figure 1B). We identified five groups of hits with distinct visual LD alterations (Figures 1B and C, Table S1): (i) six genes that, when overexpressed, resulted in a reduced number of LDs; (ii) 16 genes that induced an increase in the number of LDs; (iii) five genes that caused a strong and (iv) 14 that caused a partial clustering of LDs; and finally, (v) two genes that belonged to the desired class of factors that promoted LD accumulation in developing daughter cells. Within the latter group, overexpression of the F-box protein Rcy1, which is involved in protein recycling from endosomes to the Golgi apparatus ^46–48^, led to a moderate accumulation of LDs in daughters. Overexpression of the protein of unknown function Yer085c on the other hand caused a striking shift of the majority of cellular LDs into developing buds. Due to this strong phenotype, we set out to determine the molecular role of the poorly studied Yer085c, which we termed Lipid Droplet Motility protein 1 (Ldm1).

### The protein of unknown function Yer085c/Ldm1 promotes accumulation of LDs and mitochondria in polarized cellular regions

To re-assess the phenotype observed in the screen, we manually created a set of strains overexpressing Ldm1 from a *TEF2* promoter in combination with mCherry-fused organelle markers (Figures 2A-F). Cells were analyzed by microscopy, and the distribution of mCherry signal between mother and daughter cells was quantified. All budded cells were considered for quantification and grouped into four categories dependent on the area of the bud compared to the mother cell area (category I, 0-24%; category II, >24-39%; category III, >39-50%; category IV, >50-61%). While distribution of most organelles was not (Figures 2C and D) or only mildly (Figures 2E and F) affected by Ldm1 overexpression, LDs accumulated strongly in daughter cells across all bud size categories (Figure 2A), confirming the results of the screen. Unexpectedly, we identified mitochondria as a second organelle that responded to Ldm1 overexpression with a pronounced shift toward daughter cells (Figure 2B). We re-analyzed the set of p*TEF2*-Ldm1 strains expressing mCherry-fused organelle markers upon treatment with the mating pheromone α-factor, which induces a uniform polarization of all cells with formation of a characteristic mating protrusion. At this condition, LDs and mitochondria accumulated in the tip of the mating protrusion in response to Ldm1 overexpression, while the other organelles remained dispersed through the cell (Figure S1A), suggesting that Ldm1 does not broadly affect cell polarity or organelle transport processes, but instead specifically promotes transport of LDs and mitochondria. As a further control, we analyzed the distribution of secretory vesicles, which are an important cargo of acto-myosin-based transport to sites of bud expansion as well as to mating protrusions. Secretory vesicles were unaltered by Ldm1 overexpression both without and with α-factor treatment (Figures 2G, S1B), corroborating the notion that cell polarity and general material transport were unaffected. To obtain a high-resolution view of organelle distribution in Ldm1 overexpressing cells, we analyzed α-factor treated and vitrified Ldm1 overexpressing cells by cryo-electron tomography ^41,49^. We detected numerous LDs and mitochondria that showed a pronounced accumulation within the tip of the mating protrusion (Figure 2H).

**Figure 2.**
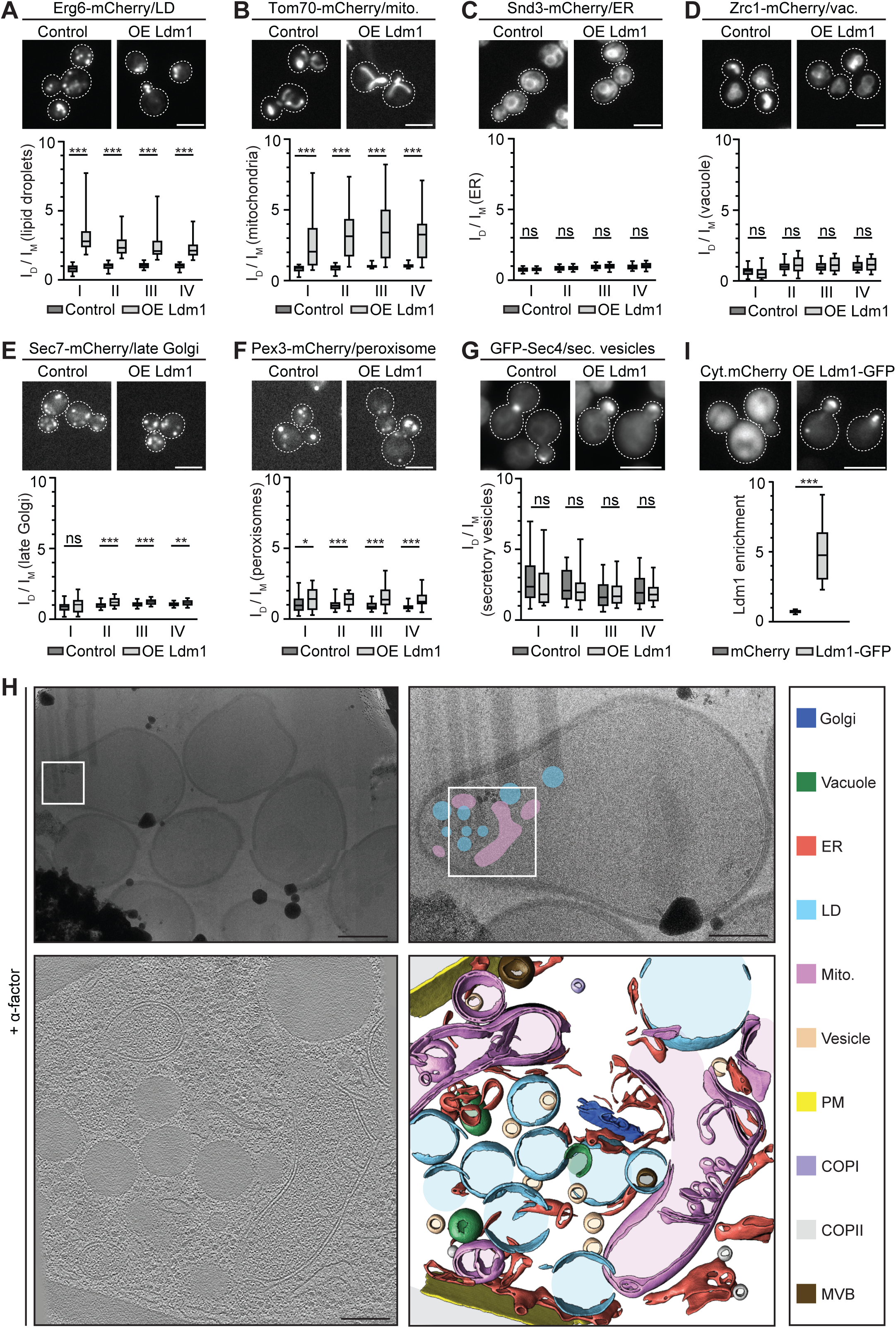
Yer085c/Ldm1 (Lipid Droplet Motility 1) overexpression results in enhanced accumulation of LDs and mitochondria at polarized cell regions. (A)-(F) Ldm1 overexpressing (*TEF2* promoter) and control cells expressing mCherry-fused organelle markers were analyzed by microscopy. Mean intensity ratios of fluorescent organelle marker signals in daughter over mother cells (I_D_/I_M_) are depicted as boxplots (5-95 percentile). Cells were categorized by bud size: I, 0-24%, II, >24-39%, III, >39-50%, IV, >50-61%. mito., mitochondria; vac., vacuole. ns, not significant; ∗, p<0.05; ∗∗, p<0.01; ∗∗∗, p<0.001. Scale bars, 5 µm. N≥60 cells, n=3. (G) Secretory vesicles were labeled by plasmid-derived GFP-Sec4 in control and p*TEF2*-Ldm1 cells and analyzed by microscopy. The ratio of the mean fluorescent signal of the vesicular structure in the bud to the mean fluorescent signal of the cytoplasm in the mother cell is shown as a boxplot (5-95 percentile). Cells were categorized by bud size: I, 0-24%, II, >24-39%, III, >39-50%, IV, >50-61%. ns, not significant. Scale bar, 5 µm. N≥60 cells, n=3. (H) Top left: transmission electron microscopy image of a FIB-milled lamella of p*TEF2*-Ldm1 cells treated for 2 hours with the mating pheromone α-factor to induce a uniform polarization of the cell population and formation of mating protrusions. Top right: zoom-in on the cell from which the tomogram was collected. LDs (blue) and mitochondria (pink) are located in the mating protrusion. Bottom left and bottom right: tomographic slice and corresponding segmentation. White squares indicate the region where the tomogram was collected. Mito., mitochondria; PM, plasma membrane; COPI, coat protein complex I; COPII, coat protein complex II; MVB, multivesicular body. Scale bars: top left, 2 µm; top right, 1 µm; bottom left, 200 nm. (I) Cells expressing Ldm1-GFP from a *TEF2* promoter were analyzed as in (A). Cells expressing cytosolic (cyt.) mCherry served as control. The ratio of the maximal intensity of fluorescence signal in the daughter cell to the maximal intensity of fluorescence signal in the cytoplasm of the mother cell is shown as a boxplot (5-95 percentile). ∗∗∗, p<0.001. Scale bar, 5 µm. N≥60 cells, n=3. Also see Figure S1.

We next wanted to assess the subcellular localization of the Ldm1 protein itself. The p*TEF2*-mCherry-Ldm1 strain from the genetic screen (Figures 1B and C) showed – despite the strong effect on LD distribution – a relatively weak mCherry signal (Figure S1C), suggesting a potential sensitivity to proteolytic loss of the tag. We created a set of Erg6-mCherry strains expressing different Ldm1 variants from the *TEF2* promoter: untagged Ldm1, N-terminally GFP-tagged Ldm1, and C-terminally GFP-tagged Ldm1. We found that the C-terminally tagged variant affected LD distribution to an extent comparable to untagged Ldm1, while N-terminal GFP tagging resulted in a markedly lower effect (Figure S1D). Both GFP-tagged Ldm1 variants localized to polarized cell regions (Figures 2I, S1D), with a lower signal intensity for the N-GFP protein (Figure S1D). The stronger effect on LDs and the stronger GFP signal suggest a preferred use of C-terminally tagged Ldm1 for detection of the protein. We generated an Ldm1-specific antiserum for western blotting, which revealed a band at the expected size of 20 kDa for untagged Ldm1, and a band migrating at 48 kDa for Ldm1-GFP, the latter one being also detected by an anti-GFP antibody (Figure S1E). Of note, we detected for the Ldm1-GFP strain a second fraction of protein migrating at the same size as untagged Ldm1 using the anti-Ldm1 antibody that was not labeled by the anti-GFP antibody (Figure S1E), again suggesting a sensitivity to proteolytic processing. For Ldm1-GFP detection by microscopy, this raises the caveat that part of the active protein is likely in an untagged state.

In summary, we have identified Ldm1 as a protein that drives accumulation of LDs and mitochondria in polarized regions of dividing as well as α-factor-treated cells.

### Ldm1 has a role in actin-based directional LD motility

As Ldm1 overexpression promotes LD accumulation in daughter cells, we assessed the effect of *LDM1* deletion on LD partitioning in dividing cells. We found that in the τι*ldm1* strain, the fraction of daughter-localized LDs was mildly reduced across all bud size classes (Figures 3A and B). Of note, while some organelles, such as mitochondria, cannot be formed de novo and thus strictly rely on being inherited from mother to daughter cells ^16,17,50^, LDs are synthesized at the membrane of the ER, indicating that bud-localized LDs can stem both from inheritance of pre-existing LDs from mothers and from motility-independent de novo LD biogenesis within daughter cells. Furthermore, LDs form physical contact sites to other organelles ^21,29^, suggesting that LDs can potentially also be dragged into buds together with partner organelles in a contact site-mediated way. We used time-resolved microscopy to compare LD motility in control, τι*ldm1*, and p*TEF2*-Ldm1 cells (Figures 3C and D). LD motility was low in τι*ldm1* cells and increased in p*TEF2*-Ldm1 cells. To test whether Ldm1-dependent LD motility was actin dependent, we incubated p*TEF2*-Ldm1 cells with latrunculin B to depolymerize actin filaments. We found that this treatment blocked Ldm1-dependent LD motility (Figure 3E). In budding or α-factor treated cells with established polarity, most LDs in p*TEF2*-Ldm1 cells were generally at any given timepoint tightly clustered around the cell pole (Figures 3C, 3E and F), partially impeding analysis of motility. We therefore attempted to create a state at which cell polarity was acutely remodeled. To achieve this, we first incubated cells with α-factor for a uniform formation of a polarized mating protrusion in all cells (Figure 3F). We then washed the cells with α-factor-free medium (α-factor release), resulting in rapid loss of polarity and selection of a new bud site distant from the mating protrusion. With this re-polarization, we observed numerous and rapid long-range motility events in p*TEF2*-Ldm1 cells (Figure 3G). Finally, to complement the data showing the latrunculin B dependence of Ldm1-induced LD motility (Figure 3E), we chose an independent genetic approach by introducing the p*TEF2*-Ldm1 allele into τι*bni1* cells which lack the main yeast formin responsible for formation of actin cables ^51,52^. *BNI1* deletion impeded Ldm1-dependent LD accumulation in daughter cells (Figures 3H and I), confirming that Ldm1 function requires an intact actin cytoskeleton. In summary, we conclude that Ldm1 promotes LD motility in a manner dependent on actin.

**Figure 3.**
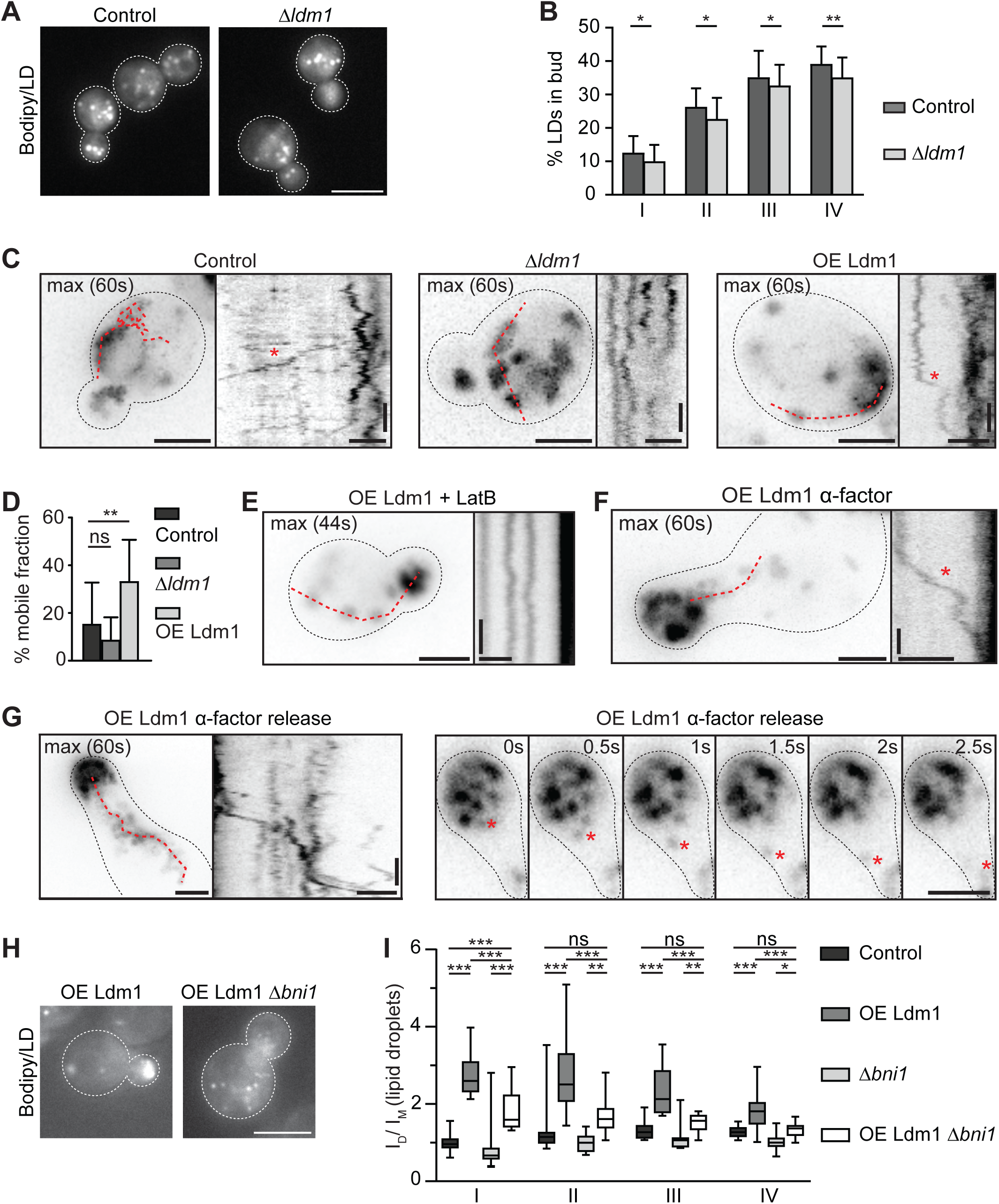
Ldm1 is involved in actin-dependent LD motility. (A) Control and *Δldm1* cells were stained with BODIPY493/503 and analyzed by fluorescence microscopy. Scale bar, 5 µm. (B) Quantification of phenotype in (A). Bar graph shows the percentage of the number of LDs in the daughter to the total number of LDs in the whole cell. Cells were categorized by bud size: I, 0-24%, II, >24-39%, III, >39-50%, IV, >50-61%. Data represented as mean ± SD. ∗, p<0.05; ∗∗, p<0.01. N≥100 cells, n=3. (C) Control, *Δldm1*, and p*TEF2*-Ldm1 (OE Ldm1) cells expressing Erg6-mCherry were analyzed by time-resolved microscopy at 2 frames per second (s). Maximal projections of 60 s and kymographs are shown. Scale bars, 2 µm. Time bars, 10 s. (D) Bar graph shows the mobile LD fraction of strains from (C). Data represented as mean ± SD. ns, not significant; ∗∗, p<0.01. n >17. (E) p*TEF2*-Ldm1 Erg6-mCherry cells were incubated with Latrunculin B for 2 hours and analyzed as in (C). Scale bars, 2 µm. Time bar, 10 s. (F) p*TEF2*-Ldm1 Erg6-mCherry cells treated with the mating pheromone α-factor were analyzed as in (C). Scale bars, 2 µm. Time bar, 10 s. (G) Left: α-factor treated p*TEF2*-Ldm1 Erg6-mCherry cells were washed three times with α-factor-free medium to release cell polarization and analyzed as in (C). Scale bars, 2 µm. Time bar, 10 s. Right: A fast LD motility event (moving LD labeled by red asterisk) is shown. Scale bar, 2µm. (H) Control and *Δbni1* cells overexpressing Ldm1-mCherry from a centromeric plasmid (*TEF2* promoter) were stained with BODIPY493/503 and analyzed by microscopy. Scale bar, 5 µm. (I) Mean intensity ratio of BODIPY493/503 signal in daughter over mother cells (I_D_/I_M_) in control and *Δbni1* cells transformed either with a plasmid for Ldm1-mCherry overexpression or empty vector is depicted as boxplots (5-95 percentile). For strains OE Ldm1 and OE Ldm1 *Δbni1*, only mCherry-positive cells were considered for quantification. ns, not significant; ∗, p<0.05; ∗∗, p<0.01; ∗∗∗, p<0.001. N=60 cells, n=3.

### Systematic approaches identify candidate Ldm1 partner proteins

We took advantage of the strong effect of *LDM1* overexpression on LD distribution to identify further genes required for LD motility via a microscopy-based screen. We used an automated mating-based assay ^53,54^ to introduce a p*TEF2*-mCherry-Ldm1 allele into a genome-wide mutant collection consisting of deletion mutants of non-essential ^55^ and hypomorphic allele mutants of essential genes ^56^. We stained the ∼6000 resulting mutant strains with the LD dye BODIPY493/503 and analyzed them by automated microscopy (Figure 4A). We identified a total of 33 genes required for efficient accumulation of LDs in daughter cells in response to *LDM1* overexpression (Figure 4B). The hits were mainly related to the actin cytoskeleton, to trafficking, and to lipid metabolism (Table S2). We complemented the genetic screen with a proteomic approach and identified physical Ldm1 partner proteins by affinity purification mass spectrometry based on stable isotope labeling with amino acids in cell culture (SILAC) ^57,58^ (Figure 4C). Among the specifically enriched proteins were Ldm1 itself, two LD surface proteins (Figure 4C, green), the translocase of the outer mitochondrial membrane (TOM complex) (Figure 4C, red), and the type V myosin motor protein Myo2 (Figure 4C, yellow). The hits of the genetic and proteomic approach served as a starting point for a mechanistic analysis of Ldm1 function (Figures 5-7 and S2-5).

**Figure 4.**
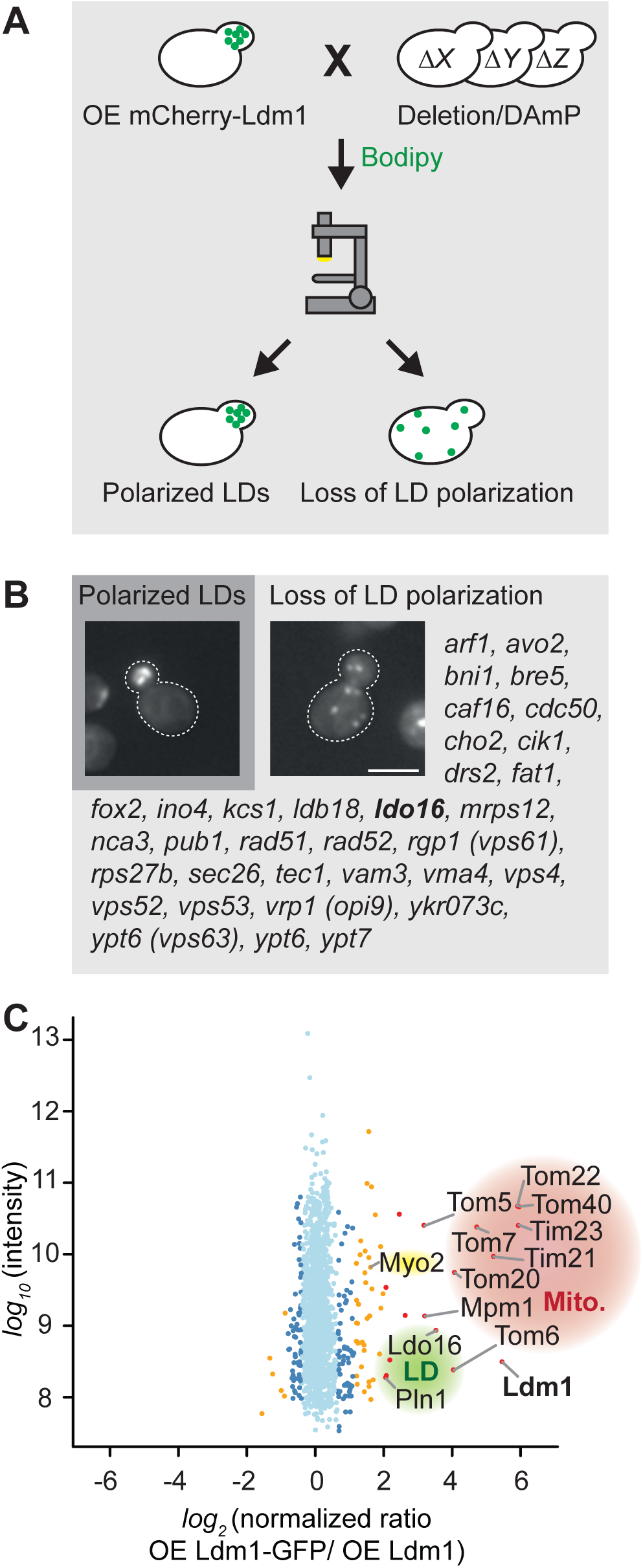
Genetic and proteomic approaches identify candidates for Ldm1 partner proteins. (A) Schematic representation of a screen designed to identify effectors of Ldm1-dependent LD accumulation at polarity sites. A p*TEF2*-mCherry-Ldm1 allele was introduced into a genome-wide collection of deletion and decreased abundance by mRNA perturbation (DAmP) mutants. Cells were stained with BODIPY493/503 and analyzed by automated microscopy. (B) Loss of function alleles that fully or partially abolish Ldm1-dependent polarized accumulation of LDs are listed. Dubious open reading frames/putative protein of unknown function genes that showed a phenotype and overlap with characterized genes are listed in brackets after the corresponding characterized gene. Gene corresponding to representative image is labeled in bold. Scale bar, 5 µm. (C) Proteomic analysis of an affinity purification of Ldm1-GFP by mass spectrometry. Analysis was performed with heavy isotope-labeled cells overexpressing Ldm1-GFP and light isotope-labeled control cells overexpressing untagged Ldm1. Protein intensities are plotted against heavy/light SILAC ratios. Significant outliers are colored in red (p<1^-11^), orange (p<1^-4^), or steel blue (p<0.05), other proteins are shown in light blue. Bait in bold, Myo2 motor protein in yellow cloud, LD proteins in green cloud, mitochondrial (mito.) proteins in red cloud. Also see Table S2.

**Figure 5.**
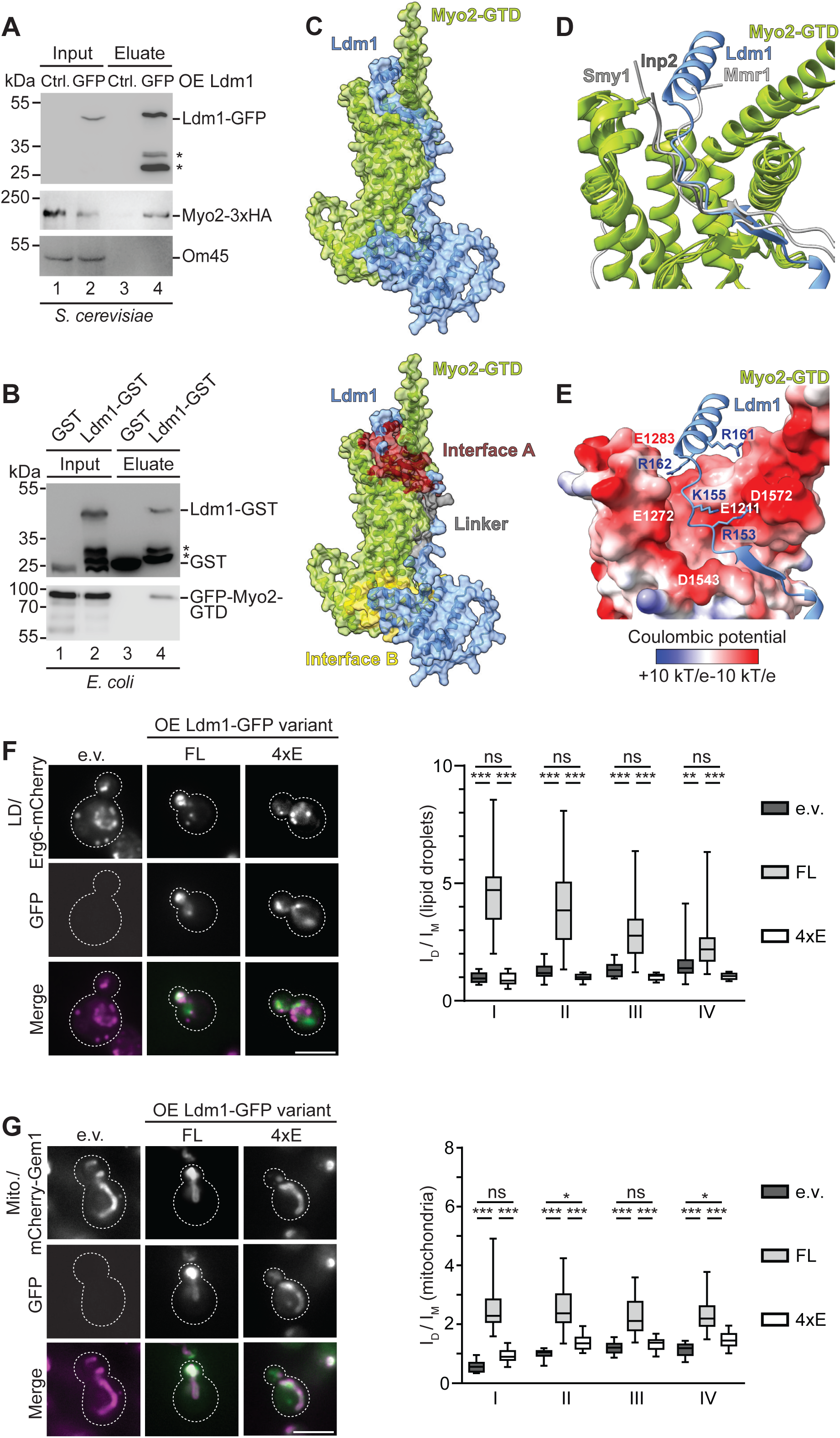
Ldm1 directly binds to the globular tail domain (GTD) of the type V myosin Myo2. (A) p*TEF2*-Ldm1-GFP cells and p*TEF2*-Ldm1 control (Ctrl.) cells expressing Myo2-3xHA were subjected to immunoprecipitation using GFP-trap, SDS-PAGE, and western blotting. Input 5%, eluate 100%. Asterisk, product of Ldm1 degradation. (B) *E. coli* cells were co-transformed with plasmids for expression of GFP-Myo2-GTD (globular tail domain) and either glutathione-S-transferase (GST) or Ldm1-GST and subjected to GST affinity purification, SDS-PAGE and western blotting. Input GST 20%, GFP 0.5%, eluate 100%. Asterisk, product of Ldm1 degradation. (C) AlphaFold 3 structure prediction of Myo2-GTD (green) in complex with Ldm1 (blue) suggesting an extended interface (ptm = 0.75, iptm = 0.71). Two potential main binding sites are highlighted in red (interface A) and yellow (interface B). (D) Close up of interface A superimposed with experimentally determined structures of Myo2-GTD (green) in complex with the Myo2 adaptor proteins Mmr1 (PDB: 6IXP, light grey) and Inp2 (PDB: 6IXR, dark grey), and the Myo2-binding kinesin related protein Smy1 (PDB: 6IXQ, grey) ^64^. (E) Close up of interface A depicting four highly conserved and positively charged arginine and lysine residues of Ldm1 (blue) that contribute to the interaction. Myo2-GTD is shown in surface representation colored by Coulomb potential and the negatively charged aspartate and glutamate residues in the interface are labelled. (F) Erg6-mCherry cells transformed with plasmids for expression of Ldm1-GFP variants (*TEF2* promoters) were analyzed by microscopy. Mean intensity ratio of fluorescent LD marker signal in daughter over mother cells (I_D_/I_M_) is depicted as boxplots (5-95 percentile). Cells were categorized by bud size: I, 0-24%, II, >24-39%, III, >39-50%, IV, >50-61%. e.v., empty vector. ns, not significant; ∗∗, p<0.01; ∗∗∗, p<0.001. Scale bar, 5 µm. N=60 cells, n=3. (G) mCherry-Gem1 cells were analyzed as in (F) to assess mitochondrial (I_D_/I_M_). e.v., empty vector. ns, not significant; ∗, p<0.05; ∗∗∗, p<0.001. Scale bar, 5 µm. N=60 cells, n=3. Also see Figures S2 and S3.

### Ldm1 physically interacts with the globular tail domain of the type V myosin Myo2

We followed up on our hits from the genetic and proteomic search for Ldm1 partner proteins (Figure 4) and first focused on the type V myosin Myo2. We tagged Myo2 with a C-terminal 3xHA tag for detection by western blot. Affinity purification of Ldm1-GFP via GFP trap resulted in an efficient co-isolation of Myo2-3xHA (Figure 5A), confirming the results of the proteomic analysis (Figure 4C).

Knowing that Ldm1 promotes actin-dependent LD motility (Figure 3) and that Ldm1 forms a complex with Myo2 (Figures 4C, 5A), we hypothesized that Ldm1 could act as a Myo2-LD adaptor protein. Myo2 adaptor proteins generally interact with the C-terminal globular tail domain (GTD) of Myo2 ^59–63^. To test experimentally if Ldm1 directly interacts with the Myo2-GTD, we co-expressed a glutathione-S-transferase (GST)-tagged Ldm1 variant and GFP-Myo2-GTD in a heterologous *E. coli* system. Purification of Ldm1 via affinity purification resulted in a co-isolation of the Myo2-GTD (Figure 5B), confirming a direct physical interaction between the two proteins and supporting the hypothesis that Ldm1 is a Myo2 adaptor. Consistently, we found that the effect of Ldm1 on LD distribution occurred independently of Myo4, the second type V myosin present in yeast (Figure S2A).

According to structure predictions, Ldm1 is a mainly α-helical protein of 173 amino acids. To map the Ldm1 binding domain that mediates the Myo2 interaction we created plasmids for expression of six Ldm1 variants lacking different regions: Ldm1_1′3-29_, Ldm1_1′30-56_, Ldm1_1′59-79_, Ldm1_1′80-124_, Ldm1_1′127-156_, and Ldm1_1′159-171_. The Ldm1 variants showed differential effects on accumulation of LDs and mitochondria in daughter cells, with Ldm1_1′3-29_ and Ldm1_1′127-156_ losing their effect on organelle distribution (Figures S2B-E). Similar results were obtained in cells treated with α-factor (Figures S3A and B).

We next used AlphaFold3 to test if an interaction surface in the Myo2-GTD can be predicted for Ldm1. A model for a complex with good confidence metrics was obtained (ipTM/interface predicted template modelling score 0.71, pTM/predicted template modelling score 0.75, with the same interaction sites predicted in all five top models) in which Ldm1 contacts Myo2-GTD extensively via two interfaces A (red) and B (yellow) (Figures 5C, S3C and D). Interestingly, the regions of Ldm1 that affected organelle distribution in our domain mapping experiment (Figures S2B-E) lie within these predicted interfaces. Amino acids 3-29 of Ldm1 form a central α-helix in the folded N-terminal domain of Ldm1 that constitutes interface B (Figures 5C, S3C and D), suggesting that deletion of this Ldm1 region will destabilize the interaction. Ldm1 amino acids 127-156 on the other hand are part of the Ldm1 region that is predicted to form interface A with the Myo2-GTD (Figures 5C, S3C and D). We compared the predicted structure of Ldm1-Myo2 with complexes between Myo2 and known Myo2-adaptors that have been determined by X-ray crystallography ^64^. We found that the Ldm1-Myo2 interface A closely resembles the structurally determined interfaces of Myo2 to the peroxisome adaptor Inp2, the mitochondrial adaptor Mmr1, and the Myo2 binding protein Smy1 ^64^ (Figure 5D). The Myo2 region at interface A forms a hydrophobic cleft lined by negatively charged residues (Figures 5E, S3E and F), while the respective region of Ldm1 contains four conserved positively charged amino acids, R153, K155, R161 and R162 (Figures 5E, S3G). To experimentally test the AlphaFold prediction, we created a variant Ldm1_4xE_ in which we swapped these four basic amino acids to glutamates to disrupt interface A. This mutation resulted in a loss of the Ldm1 effects on LD and mitochondrial distribution (Figures 5F and G). In summary, we find that Ldm1 directly interacts with the Myo2-GTD and provide evidence supporting an Ldm1-Myo2 binding mode similar to known Myo2 adaptor proteins.

### Ldo16 is the LD surface receptor for Ldm1

Ldm1 promotes LD motility and physically interacts with Myo2, indicating that it acts as Myo2-LD adaptor protein. However, according to structure predictions, Ldm1 lacks hydrophobic domains for direct LD binding. We therefore asked how binding of Myo2-Ldm1 complexes to LDs is achieved. In our proteomic search for Ldm1 partner proteins (Figure 4C), two LD proteins were identified that could act as LD surface receptors for the Myo2-Ldm1 complex. One of them is the structural LD protein Pln1 ^65^, which was moderately enriched with Ldm1 (normalized heavy/light ratio: 4.2). The second one was Ldo16, a multifunctional LD protein that interacts with the seipin LD biogenesis machinery ^38–40^ and has a role in formation of the vacuole lipid droplet contact site vCLIP together with the vacuole surface protein Vac8 ^41,42^. Ldo16 was efficiently co-enriched with Ldm1 in the proteomics experiment (normalized heavy/light ratio: 11.5), and additionally was a hit in the genetic screen for factors required for Ldm1-dependent LD accumulation in daughter cells (Figures 4A and B, Table S2).

Ldm1-GFP trap and western blot confirmed that Ldo16 was efficiently co-isolated with Ldm1, while Pln1 was only mildly co-enriched (Figure 6A). We manually re-created p*TEF2*-Ldm1 τι*ldo16* and p*TEF2*-Ldm1 τι*pln1* strains and analyzed them by BODIPY493/503 staining and microscopy. While loss of Pln1 did not affect the p*TEF2*-Ldm1 phenotype (Figure S4A), loss of Ldo16 abolished the ability of Ldm1 to induce LD accumulation in buds during cell division or in mating protrusions of cells treated with α-factor (Figures 6B and C, S4A and B). Ldo16 re-expression from a plasmid fully restored the Ldm1 phenotype (Figures 6B and C, Figure S4B), consistent with a role of Ldo16 as Ldm1 receptor. Time-resolved microscopy showed that *LDO16* deletion blocked Ldm1-dependent LD motility (Figure 6D), corroborating the involvement of Ldo16 in the LD motility machinery.

**Figure 6.**
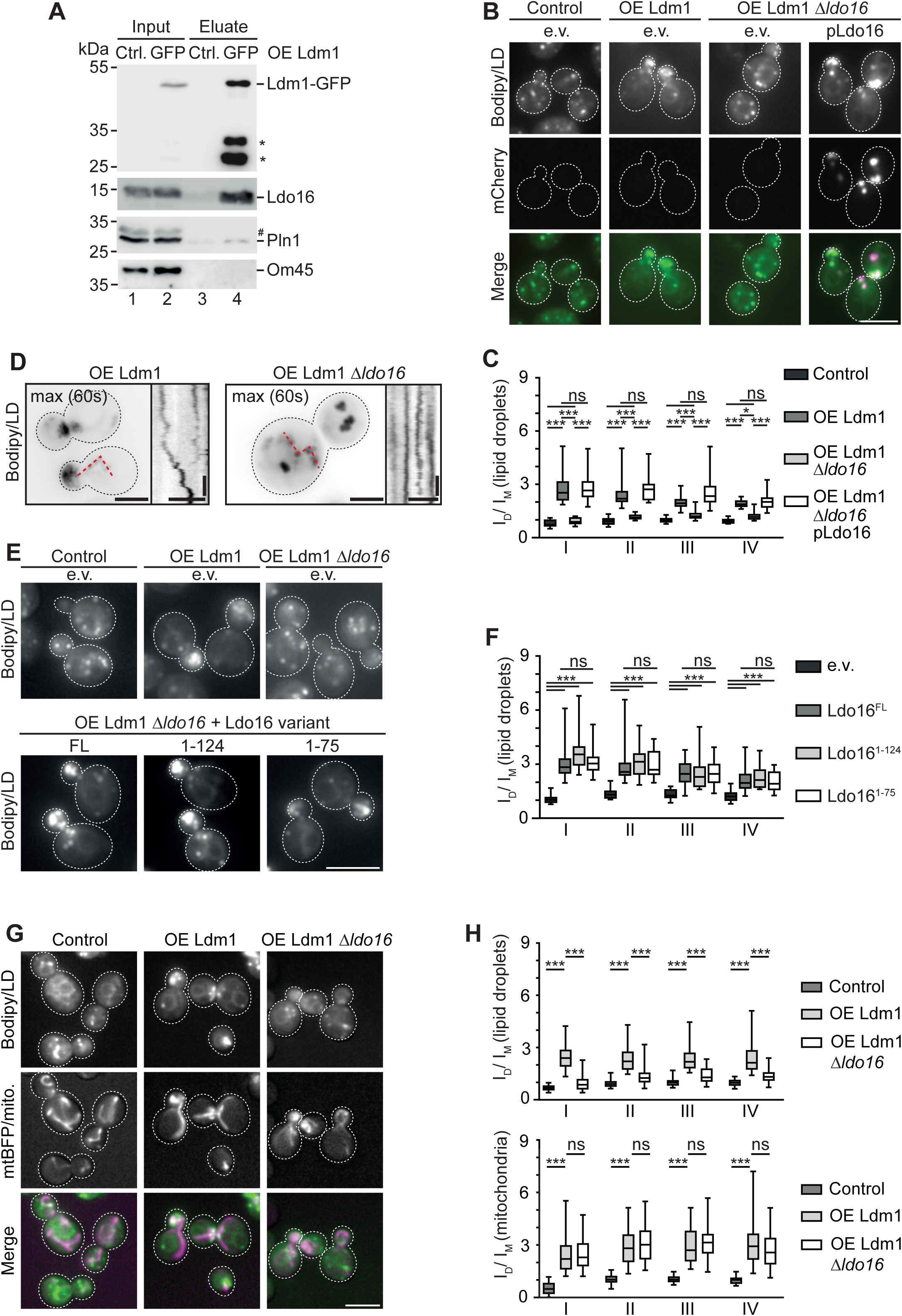
The multifunctional LD surface receptor Ldo16 is required for Ldm1-dependent LD motility, but dispensable for mitochondrial transport. (A) p*TEF2*-Ldm1-GFP cells and p*TEF2*-Ldm1 control (Ctrl.) cells were subjected to immunoprecipitation using GFP-trap, SDS-PAGE, and western blotting. Input 5%, eluate 100%. Asterisk, product of Ldm1 degradation. #, unspecific band. (B) Control, p*TEF2*-Ldm1, and p*TEF2*-Ldm1 *Δldo16* cells were transformed with indicated plasmids and stained with BODIPY493/503. pLdo16, centromeric plasmid for expression of Ldo16-mCherry from the p*LDO16* promoter. e.v., empty vector. Scale bar, 5 µm. (C) Mean intensity ratio of fluorescent BODIPY493/503 signal in daughter over mother cells (I_D_/I_M_) is depicted as boxplots (5-95 percentile). Cells were categorized by bud size: I, 0-24%, II, >24-39%, III, >39-50%, IV, >50-61%. ns, not significant; ∗, p<0.05; ∗∗∗, p<0.001. Scale bar, 5 µm. N=60 cells, n=3. (D) p*TEF2*-Ldm1 cells and p*TEF2*-Ldm1 *Δldo16* cells were labeled with BODIPY493/503 and analyzed by time-resolved microscopy at 2 frames per seconds (s). Maximal projections of 60 s, and kymographs are shown. Scale bars, 2 µm: Time bars, 10 s. (E) Control, p*TEF2*-Ldm1, and p*TEF2*-Ldm1 *Δldo16* cells were transformed with indicated plasmids and stained with BODIPY493/503. FL, centromeric plasmid for expression of full length Ldo16-mCherry from the p*LDO16* promoter. 1-124 and 1-75, centromeric plasmids for expression of truncated Ldo16 variants comprising amino acids 1-124 or 1-75 from the p*LDO16* promoter. e.v., empty vector. Scale bar, 5 µm. (F) Mean intensity ratios of fluorescent BODIPY493/503 signals in daughter over mother cells (I_D_/I_M_) are depicted as boxplots (5-95 percentile). Cells were categorized by bud size: I, 0-24%, II, >24-39%, III, >39-50%, IV, >50-61%. ns, not significant; ∗∗∗, p<0.001. Scale bar, 5 µm. N=60 cells, n=3. (G) Control, p*TEF2*-Ldm1, and p*TEF2*-Ldm1 *Δldo16* cells were transformed with a plasmid for expression of mtBFP to label mitochondria and stained with BODIPY493/503. Scale bar, 5 µm. (H) Mean intensity ratios of fluorescent organelle signals in daughter over mother cells (I_D_/I_M_) are depicted as boxplots (5-95 percentile). Cells were categorized by bud size: I, 0-24%, II, >24-39%, III, >39-50%, IV, >50-61%. ns, not significant; ∗∗∗, p<0.001. Scale bar, 5 µm. N≥60 cells, n=3. Also see Figure S4.

We performed a structure-function analysis to determine the Ldo16 subdomain required for Ldm1-dependent LD motility. Ldo16 has a total length of 148 amino acids, with the N-terminal 55 amino acids forming a hydrophobic, LD integral domain, and the C-terminal 93 amino acids being exposed to the cytosol ^41,42^. The N-terminal hydrophobic domain as well as the adjacent amino acids 56-75 are critically required for correct Ldo16 targeting to LDs ^41,42^. We used two Ldo16 truncation constructs, Ldo16_1-124_ and Ldo16_1-75_, that have previously been used to map the interaction of Ldo16 with its partner protein Vac8 at vCLIP contact sites ^41^. We found that both truncation variants Ldo16_1-124_ and Ldo16_1-75_ efficiently supported Ldm1-dependent LD accumulation in daughters (Figures 6E and F) and mating protrusions (Figure S4C) efficiently. The latter Ldo16 variant exposes only around 20 amino acids to the cytosol, suggesting that this region of the protein is key for the interplay with Ldm1. Of note, variant Ldo16_1-75_ is defective in formation of vCLIP ^41^, indicating that Ldm1- and Vac8 binding depend on distinct Ldo16 subdomains.

We asked if Ldo16 was also involved in Ldm1 effects on mitochondrial distribution. We therefore simultaneously monitored LDs using BODIPY493/503 and mitochondria using mtBFP in p*TEF2*-Ldm1 and p*TEF2*-Ldm1 τι*ldo16* cells. We found that while Ldm1-dependent LD accumulation in buds and mating protrusions was blocked by *LDO16* deletion, mitochondrial accumulation was unaffected (Figures 6G and H, S4D), showing that *LDO16* deletion allows for a genetic separation of the two Ldm1-dependent effects. In summary, we have identified Ldo16 as the LD surface receptor for Ldm1 that is specifically required for motility of LDs, but dispensable for mitochondria, and have mapped a minimal Ldo16 region sufficient for Ldm1 binding.

### Ldm1 overexpression rescues mutants of the Myo2 adaptors Mmr1/Ypt11

Finally, we assessed the link of Ldm1 to the translocase of the outer mitochondrial membrane TOM (Figure 4C). Western blot confirmed a co-isolation of the subunits of this essential protein translocase together with Ldm1-GFP via GFP trap (Figure 7A). Due to this link of Ldm1 to a mitochondrial outer membrane complex and the observation that Ldm1 overexpression promotes accumulation of mitochondria in daughter cells (Figures 2B, 5G, 6G and H, S2D and E) and mating protrusions (Figures 2H, S1A, S3B, S4D), we hypothesized that Ldm1 might act as Myo2-mitochondria adaptor in addition to its role as Myo2-LD adaptor. We analyzed mitochondrial partitioning between mothers and daughters in τι*ldm1* cells and found that unlike for LDs (Figures 3A and B), the bud-localized mitochondrial fraction was unaffected (Figure 7B). This is not surprising as two mitochondrial Myo2 adaptors have already been identified, Mmr1 ^17^ and Ypt11 ^18^, indicating that redundant machineries are in place for mitochondrial motility. To test whether Ldm1 acts in parallel to the known Myo2-mitochondria adaptors or whether its effect depends on these proteins, we used two strains τι*ypt11 mmr1-20* and τι*ypt11 mmr1-5*, which both carry a *YPT11* deletion and a temperature-sensitive *MMR1* allele ^16^. We analyzed these cells at the permissive temperature of 26°C and the restrictive temperature of 37°C and observed mitochondria using mtBFP and LDs using BODIPY493/503 (Figures 7C-E, S5A-E). Consistent with previous reports ^16^, we found that at the restrictive temperature, daughter cells were virtually completely devoid of mitochondria, while LDs were regularly distributed (Figures 7D and E, S5D and E). It has previously been shown that the mitochondrial transport defect can be rescued via a plasmid that encodes a Myo2 variant that is synthetically targeted to the mitochondrial surface via fusion with the outer membrane protein Fis1 ^16,66^. We reproduced this experiment and compared the effect of the Myo2-Fis1 plasmid to overexpression of Ldm1. We found that Ldm1 restored mitochondrial transport to buds similarly to Myo2-Fis1 (Figures 7D and E, S5D and E). This shows that Ldm1 does not require Mmr1 or Ypt11 for its effect on mitochondria and is consistent with a role of Ldm1 as a third Myo2 adaptor protein. Of note, while Ldm1 promoted accumulation of both mitochondria and LDs in daughter cells as expected, Myo2-Fis only promoted transport of mitochondria, while LDs were unaffected (Figures 7D, S5D), supporting the notion that Ldm1 is a bifunctional Myo2-adaptor.

**Figure 7.**
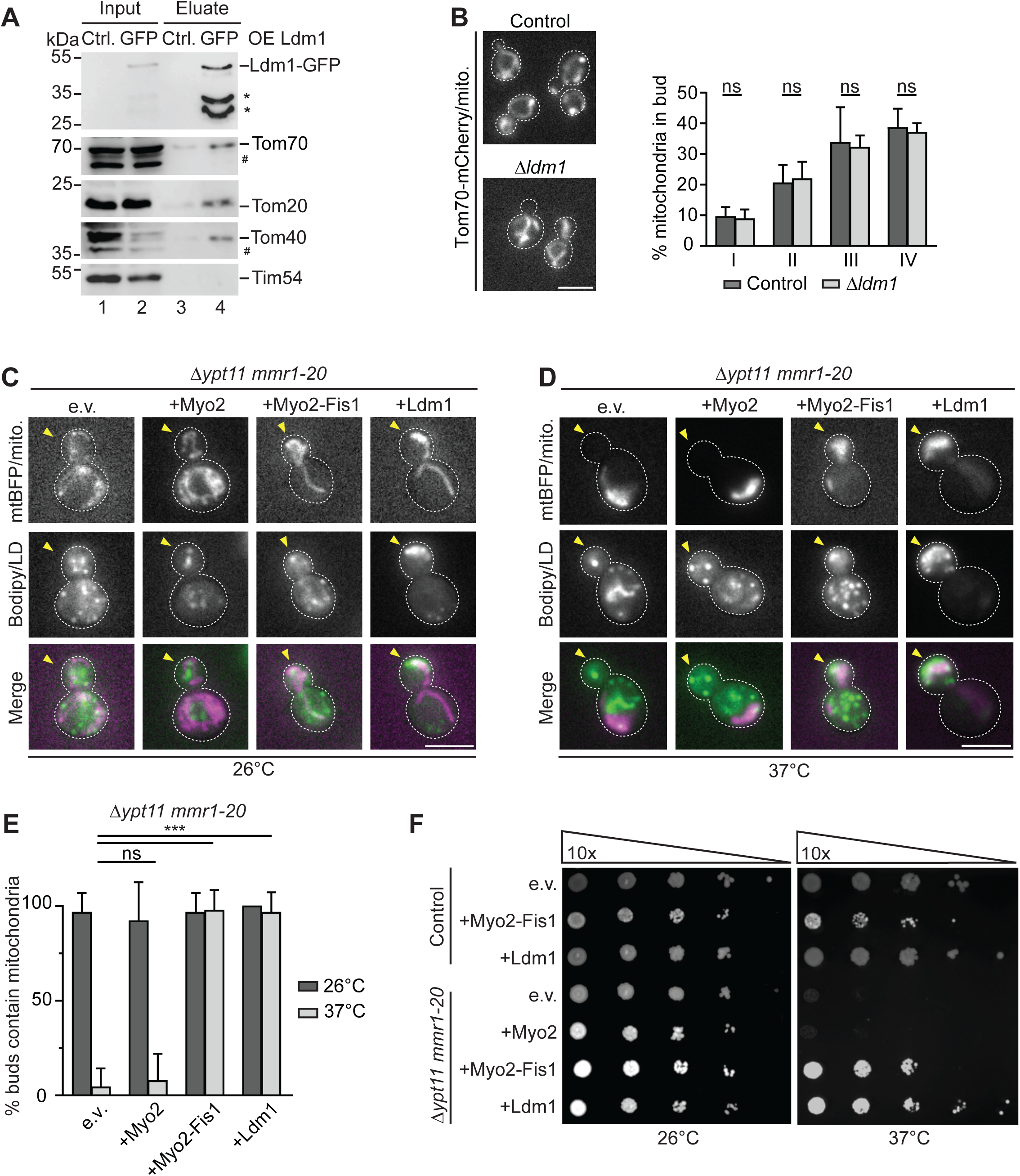
Ldm1 overexpression rescues mitochondrial transport and viability of mutants defective in the mitochondrial Myo2 adaptors Ypt11 and Mmr1. (A) p*TEF2*-Ldm1-GFP cells and p*TEF2*-Ldm1 control (Ctrl.) cells were subjected to immunoprecipitation using GFP-trap, SDS-PAGE, and western blotting. Input 5%, eluate 100%. Asterisk, product of Ldm1 degradation. #, unspecific bands. (B) Control and *Δldm1* cells expressing Tom70-mCherry to label mitochondria were analyzed by microscopy. Percentage of signal thresholded mitochondrial area in daughter cells over total mitochondrial area was determined. Data represented as mean ± SD. Scale bar, 5 µm. N≥120 cells, n=3. (C) *Δypt11 mmr1-20* cells expressing mtBFP for labeling of mitochondria were transformed with plasmids for overexpression of Myo2 (+Myo2), a Myo2 variant fused to the outer mitochondrial membrane protein Fis1 (+Myo2-Fis1) for attachment of the motor protein to mitochondria independent of Myo2-adaptor proteins, or Ldm1 (+Ldm1). Cells were cultured at 26°C, stained with BODIPY493/503, and analyzed by microscopy. e.v., empty vector. Scale bar, 5 µm. (D) Same as in (C), with the difference that cells were incubated at 37°C for 1.5 hours prior to and during imaging. (E) The percentage of dividing cells with mitochondria present in the bud was determined. Data represented as mean ± SD. ns, not significant; ∗∗∗, p<0.001. N≥60 cells, n=3. (F) Control and *Δypt11 mmr1-20* cells transformed with plasmids for overexpression of Myo2 (+Myo2), a Myo2 variant fused to the outer mitochondrial membrane protein Fis1 (+Myo2-Fis1) for attachment of the motor protein to mitochondria independent of Myo2-adaptor proteins, or Ldm1 (+Ldm1) were grown overnight at 26°C, adjusted to an OD_600_ of 0.05, serially 10-fold diluted, spotted on agar plates and grown at 26°C or 37°C for 3 days. e.v., empty vector. Also see Figure S5.

Mitochondria cannot be formed de novo, and they fulfill essential functions. Defects in mitochondrial inheritance therefore result in severely sick ^50^ or lethal phenotypes ^16,17^. We assessed viability of τι*ypt11 mmr1-20* and τι*ypt11 mmr1-5* cells using a drop assay. While growth of these cells was blocked at the non-permissive temperature, viability was restored by expression of Myo2-Fis as reported (Figures 7F, S5F) ^16^. Furthermore, we found that overexpression of Ldm1 efficiently rescued growth of both *ypt11 mmr1-ts* mutants (Figures 7F, S5F).

In conclusion, Ldm1 can restore mitochondrial inheritance and cell viability in *ypt11 mmr1* mutants, corroborating its role as Myo2-organelle adaptor for LDs and mitochondria.

## DISCUSSION

We have identified the protein of unknown function Yer085c, which we term Lipid Droplet Motility 1 (Ldm1), as a Myo2 adaptor protein that mediates actin-dependent LD transport together with its receptor Ldo16. Additionally, Ldm1 affects mitochondrial motility independent of the known mitochondrial Myo2 adaptors Mmr1 and Ypt11.

Elevated Ldm1 levels promote LD motility and LD accumulation in developing daughter cells, while loss of the protein results in decreased motility. Ldm1-dependent effects on LDs are inhibited upon latrunculin B treatment and in a *bni1* mutant, showing that they depend on a functional actin cytoskeleton. Ldm1 forms a complex with the type V myosin Myo2 in vivo. Direct binding of Ldm1 to the Myo2 GTD upon heterologous expression, structure predictions, and mutational analyses suggest that Ldm1-Myo2 interaction occurs at the Myo2 region that also binds to the Myo2 adaptor proteins Inp2 and Mmr1 and the Myo2 interactor Smy1 ^64^.

LD motility is a wide-spread phenomenon, and the question why LDs move is heavily debated ^20,21^. Yeast cells undergo an asymmetric cell division, necessitating active delivery of organelles from mother to daughter cells ^67^. While active inheritance is strictly required for organelles that cannot be generated de novo, such as mitochondria, LDs can be formed from the ER, so that LD inheritance is not crucial. Nonetheless, LD inheritance has been described ^10,37^. It is conceivable that delivery of lipid-packed LDs might be beneficial for the efficient development of growing daughter cells, by providing substrates for energy metabolism and building blocks for the growing membranes. In this respect, LDs might be a very good source of sterols, which can be liberated from stored sterol esters by sterol ester hydrolases. Large amounts of sterols are required at the growing plasma membrane of the daughter cell, which has a particularly high sterol concentration. A role of LD motility for ensuring nutrient supply has also been discussed in other organisms, for example in developing fly embryos, which show a high degree of LD motility ^68,69^. In the fly embryo, it has also been discussed that LD motility could have a role in efficient clearance of toxic lipid or protein species from all parts of the cell ^70^. In mammalian cells, an interplay of LDs with cytoskeletal elements has been related to the control of LD formation ^35^ or LD turnover by modulating LD-LD clustering and thus access of lipases to the LD surface ^33^. Moreover, LD motility modulates triglyceride secretion from hepatocytes in response to changes in feeding state ^36^. Finally, there are indications that LD motility likely is linked to the reorganization of the LD contact site landscape. Due to their unique phospholipid monolayer architecture, LDs do not participate in organelle communication via bulk trafficking of bilayer vesicles. Instead, they appear to rely heavily on communication via physical contact sites ^22,29–31^. LD formation and growth depend on contact sites with the ER, while contacts to other LD partner organelles, such as mitochondria, peroxisomes, and lysosomes/vacuoles, can promote LD breakdown. It is conceivable that strategic reorganizations of LD contact sites depend on LD motility, and indeed, alterations of the cytoskeleton are linked to alterations in LD contact sites ^32,34^. The latter proposed function of LD motility is particularly interesting in light of our finding that Ldm1 binds to LDs via an interaction with the LD surface protein Ldo16, which is additionally involved in LD contact sites. Yeast Ldo16 as well as its putative human homolog LDAF1/promethin and the fly homolog dmLDAF1 cooperate with the seipin machinery ^38–40,71–73^. Seipin has a key role in the formation of LDs from the ER, and its loss results in the lipid storage disease lipodystrophy ^74^. After LD maturation, seipin stays localized to LD-ER contact sites where it plays a role in LD maintenance and communication with the ER ^75,76^. A further partner protein of Ldo16 is the vacuole surface protein Vac8, and Ldo16-Vac8 complexes act as tethers for formation of vacuole lipid droplet (vCLIP) contact sites. vCLIP contacts are formed predominantly at nutrient restriction, and are required for efficient lipophagy, a selective form of LD autophagy ^41,42^. Thus, Ldo16 is a multifunctional LD surface receptor protein that has at least three different roles in LD biology, (i) together with seipin at LD-ER contact sites and in LD formation, (ii) together with Vac8 at LD-vacuole contact sites and in lipophagy and (iii) together with Ldm1 in LD motility. This suggests a mechanistic coordination between LD confinement to specific partner organelles by local physical tethering, and LD coupling to the motility machinery, a link that may likely offer functional benefits for the cell. For instance, it is feasible that LD motility requires an uncoupling from organelle contact sites. Ldo16 could thus have a role in mediating a switch from contact site-based LD retention to Ldm1-based LD motility. An important task for the future will be to describe in detail how the different functions of Ldo16 are related to the metabolic state of the cell. Interestingly, shared machineries for contact sites and motility have also been observed in other systems. Vacuole motility depends on the Myo2 adaptor Vac17 and the vacuole surface receptor Vac8 ^14,15^. The latter protein also acts as vacuole-nuclear ER tether ^77^, and as vCLIP tether protein ^41,42^. Thus, the phenomenon of a direct structural link between organelle contact sites and organelle motility is shared between LDs and vacuoles.

In addition to its role in LD motility, we found that Ldm1 also promotes motility of mitochondria, an observation that opens exciting perspectives for future studies. In the past, two mitochondrial Myo2 adaptor proteins have been described, Ypt11 ^18^ and Mmr1 ^17^. We find that Ldm1 overexpression rescues mitochondrial inheritance and viability of *ypt11 mmr1* mutants, indicating that Ldm1 acts in a parallel pathway, and likely represents a third mitochondrial Myo2 adaptor. Uncovering in the future physiological or stress conditions that differentially affect the adaptors may help to tease apart the exact roles of these seemingly redundant proteins. Affinity chromatography suggests that Ldm1 binding to the mitochondrial surface could be mediated by an interaction with the TOM complex, an essential protein translocase of the outer mitochondrial membrane. A more detailed analysis will be required to determine the mechanistic basis of Ldm1 binding, and to address a possible link to protein translocation. Of note, TOM subunits have been found to be involved in mitochondrial contact sites to the ER and vacuoles ^78,80,82,84,86^, suggesting that Ldm1 could possibly link contact site formation and motility for both lipid droplets and mitochondria. Ldm1-dependent mitochondrial motility can be genetically uncoupled from LD motility via *LDO16* deletion. Future studies will hopefully uncover how Ldm1-dependent effects on mitochondria and LDs are interrelated on a mechanistic and on a functional level.

## ACKNOWLEDGEMENTS

This work was supported by the Gerty Cori Programme, Medical Faculty, University of Münster (to M.B.), and the Deutsche Forschungsgemeinschaft (DFG, German Research Foundation), SFB1190 P21 (to M.B.) and P22 (to R.F.-B), SFB1348 A13 (to M.B.) and A01 (to R.W.S.), SFB1557 P03 (to M.B.), P06 and Z1 (to F.F.), P10 (to D.K.) and P15 (to R.W.S.), and FOR5815 P6 (to M.B.). F.F. acknowledges funding from the Heisenberg programme of the DFG (491484150). R.F.-B. acknowledges funding from Germany’s Excellence Strategy (EXC 2067/1-390729940). Cryo-electron tomography instrumentation at the University of Göttingen was jointly funded by the DFG Major Research Instrumentation program (448415290) and the Ministry of Science and Culture of the State of Lower Saxony. D.T.V.D. and M.H. are members of CiM-IMPRS, the joint graduate school of the Cells-in-Motion Interfaculty Centre, University of Münster, Germany and the International Max Planck Research School – Molecular Biomedicine, Münster, Germany. We thank Anthony Bretscher, Maya Schuldiner, Tommer Ravid and Naama Barkai for strains and plasmids, Christoph Thiele for providing LD540, Nikolaus Pfanner for antibodies, and all members of the Bohnert lab for discussions.

## AUTHOR CONTRIBUTIONS

Conceptualization: M.B.; Investigation and Formal Analysis: X.-T.Z., D.T.V.D, L.P., R.M.F., M.H., P.H., B.E., J.C., S.W., M.W., C.S., R.W.S., MB; Visualization: X.-T.Z., D.T.V.D, L.P., R.M.F., M.H., P.H., B.E., J.C., S.W., M.W., C.S., R.W.S., MB; Writing – Original Draft: M.B.; Writing – Review and editing X.-T.Z., D.T.V.D, L.P., R.M.F., M.H., P.H., B.E., J.C., S.W., M.W., D.K., C.S., R.F.-B., F.F., R.W.S., MB; Supervision and Funding Acquisition: M.B., R.W.S., F.F., R.F.-B., D.K.

## DECLARATION OF INTERESTS

The authors declare no competing interests.

## MATERIALS AND METHODS

### Yeast strain backgrounds

Yeast strains used in this study are derivatives of the *Saccharomyces cerevisiae* BY4741 (*MATa his3Δ1 leu2Δ0 met15Δ0 ura3Δ0*) ^79^ or a related wild-type that carries mutations required for creation of genome-wide mutant collections via automated mating (*MATα his3Δ1 leu2Δ0* LYS2+ *met15Δ0 ura3Δ0 can1Δ::STE2pr-spHIS5 lyp1Δ::STE3pr-LEU2*) ^56^.

### Yeast cell culture

Yeast cells were grown overnight in synthetic medium (0.67% weight/volume yeast nitrogen base with ammonium sulfate, amino acid supplements: alanine, arginine, asparagine, aspartic acid, cysteine, glutamine, glutamic acid, glycine, inositol, isoleucine, lysin, methionine, para-aminobenzoic acid, phenylalanine, proline, serine, threonine, tyrosine, valine, histidine, leucine, uracil, tryptophan, adenine) supplemented with 2% glucose at 30°C in a shaking incubator at 280 rpm. Subsequently, cells were diluted and grown until they reached the logarithmic growth phase. For experiments using temperature-sensitive strains, cells were grown at 26°C to reach the logarithmic phase, subsequently cells were shifted to 37°C for 1.5-2 hours. For α-factor treatment, MATa cells were incubated with α-factor at a final concentration of 10 µg/mL at 30°C for 30 minutes. For α-factor release, after α-factor treatment for 2 hours, cells were washed with α-factor-free synthetic medium three times.

For Latrunculin B treatment, cells were exposed to a final concentration of 200 µM Latrunculin B at 30°C for 2 hours.

### Generation of plasmids and yeast strains

Plasmids were generated by the Gibson assembly cloning method. PCR products and digested vectors were mixed with NEBuilder HiFi DNA Assembly Master Mix (NEB) and incubated at 50°C for 15 minutes. Reactions were briefly cooled down and transformed into competent *Escherichia coli* cells. For creation of plasmid Ldm1(4xE) carrying four point mutations (glutamates at positions 153, 155, 161, 162), a one-step multiple-site plasmid mutagenesis method was used ^81^. Primers for PCR-based genomic manipulation of yeast strains via homologous recombination were designed through a webtool, Primers-4-Yeast (https://www.weizmann.ac.il/Primers-4-Yeast/) ^83^. Yeast transformations were conducted with a Li-acetate/PEG mix (100 mM lithium acetate; 10 mM Tris-HCl (pH 8), 1 mM EDTA, 40 % (w/v) polyethylene glycol 3350), carrier DNA, and plasmid or PCR-derived DNA ^85^.

### Yeast cell growth assay

Yeast cells were pre-cultured overnight in synthetic medium supplemented with 2% glucose at the indicated temperature. On the next day, cells were diluted and grown until reaching an OD_600_ around 1. Cell concentration was adjusted to an OD_600_ of 0.05, cells were serially 10-fold diluted, 5 µL of each dilution was spotted on the indicated plates and grown at indicated temperature for up to 3 days.

### Automated library preparation and handling

Query strain p*TEF2*-mCherry-Ldm1 from the SWAT p*TEF2*-mCherry mutant collection ^44,45^ carrying mutations required for automated mating and mutant selection was crossed with the genome-wide deletion ^55^ and DAmP ^56^ mutant collections. Double mutant strains were obtained through mating, diploid selection, sporulation induction (7 days), haploid selection and final mutant selection ^53,54^. A subset of strains from the resulting new mutant collection was validated by check PCR, linkage analysis and microscopy. The mutant collection was handled with a RoToR bench-top colony array instrument (Singer instruments).

### Fluorescence microscopy and high-throughput microscopy

For high-content imaging, cells were grown to logarithmic phase in 384 well polysterene plates and stained with BODIPY493/503, then moved to 384 well glass bottom plates (Brooks) coated with Concanavalin A (Sigma-Aldrich) using the Bench Smart 96 liquid handler (Mettler Toledo). For analysis of small sample numbers, cells were grown to logarithmic phase in glass tubes and manually moved to a Concanavalin A coated glass-bottom microscope plate. After 15 minutes, wells were washed once with respective medium to maintain adherent cells. For LD staining, cells were incubated with BODIPY493/503 (1 µM; Sigma-Aldrich) for 15 minutes, or with LD540 (0.5 µg/mL) for 30 minutes, and washed once with medium. Subsequently, cells were imaged at room temperature or the indicated temperature with the Olympus screening station ScanR. Images were acquired with the Olympus ScanR Automated Image Acquisition Software using an Olympus IX83 inverted fluorescence microscope with a Lumencor SpectraX LED light source and a 40x or 60x air objective and reviewed using the ImageJ analysis program. For tracking of LD motility (Figures 1A, 3C-G, 6D), epifluorescence time series were acquired on an iMIC-based microscope (FEI/Till Photonics) equipped with Olympus 100x/1.49 NA oil immersion objective and a Sole-5 laser box (405/477/561/640 nm, Omikron). Images were captured with an iXon Ultra 897 EMCCD (Andor) cooled to -80 °C. Unless stated otherwise, all images were acquired with an electron gain of 300, preamplifier gain of 1 and no binning. Acquisition was controlled by the LiveAcquisition software (FEI/Till Photonics). For analysis of LD motility (Figures 1A, 3C-G, 6D) images were processed in ImageJ. Time series were bleach-corrected using an exponential fit. Maximum temporal projections over the indicated time were generated with the “Z-project” command. Kymographs were generated along the indicated dotted lines using the “Reslice” command. Images are shown in inverted grayscale. Images were contrast adjusted and scaled in the final figures for illustration purposes only.

### Cryo-electron tomography

Sample vitrification: α-factor treated cells overexpressing Ldm1 were cultured to OD_600_ 0.8. A drop of 3.5 µL of culture was deposited on the carbon side glow-discharged Cryo-EM grids (R1.2/1.3, Cu 200 mesh grid, Quantifoil microtools) mounted on a Vitrobot Mark IV (Thermo Fisher Scientific). Grids were back-blotted with filter paper (Whatman 597) to remove the excess liquid prior to vitrification by quick plunging into a liquid ethane/propane mixture at liquid nitrogen temperature, and stored in grid boxes in liquid nitrogen until further need.

Sample thinning: An Aquilos 2 Cryo-focused ion beam (FIB)/scanning electron microscope (SEM) (Thermo Fisher Scientific) was used to prepare lamellae. Organometallic platinum was deposited on the grid with the gas injection system (GIS) for 40 seconds to protect the milled region from the ion beam. The sample was tilted to an angle of 20° and 200 nm thick lamellae were prepared in two steps: first, using the Ga2+ ion beam at 30 kV and 300 pA beam current for rough milling and then by fine milling at 30 kV and 50 pA. The milling process was monitored using SEM imaging at 3 kV and 13 pA.

Data acquisition: Data collection was performed using a Krios G4 Cryo-transmission electron microscope (Thermo Fisher Scientific) equipped with a 300kV field emission gun, Selectris energy filter, and a Falcon 4i direct electron detector camera, at a magnification of 33000x (3.653 Å/pixel) at a defocus of -5 to -7 µm. Tilt series from - 46° to +64° at increments of 3° of the lamellae were acquired using the dose-symmetric acquisition scheme ^87^ in SerialEM ^88^ and a target total dose per tomogram around 120 e-/A2. Using the camera in dose-fractionation mode, between 700 to 900 EER frames were generated per tilt image. MotionCor2 ^89^ was used to align the frames and the new tilts series were reconstructed using patch tracking in IMOD 4.11 ^90^ and weighted back-projected for tomogram reconstruction. Tomograms were binned to 14.61 Å/pixel and filtered using a Wiener-like filter implemented in the MATLAB script tom_deconv ^91^.^91^

Tomogram segmentation: automatic membrane segmentation was performed using MemBrain-Seg ^92^. The segmentation was manually corrected and colored using Amira (Thermo Fisher Scientific). Images were produced using UCSF Chimera ^93^.

### Protein extraction, SDS-PAGE, western blotting

Cells were grown to logarithmic growth phase and an equivalent of 2.5 OD_600_ was harvested by centrifugation. Proteins were extracted through alkaline lysis ^94^ and were analyzed by SDS-PAGE and western blotting. Primary and secondary antibodies used in this study are listed in the key source table. The antibody signal was detected by enhanced chemiluminescence (ECL) using an Azure Imaging system.

### GFP immunoprecipitation

Equal amounts of cells expressing indicated proteins fused with a GFP tag and controls were grown to OD_600_ 0.8, harvested from cultures, washed with distilled water, and collected via centrifugation. Cell pellets were rapidly frozen in liquid nitrogen. Samples were resuspended in 500 µL GFP pulldown buffer (150 mM KOAc, 20 mM HEPES pH 7.4, 5% glycerol, cOmplete protease inhibitor and 1% Octyl-beta-Glucoside (Thermo Fisher)) and 500 µL glass beads (Sigma-Aldrich) were added. Cells were lysed using a shaker (IKA VIBRAX). 50 µL cell lysates of each sample were collected as “input”, mixed with HU buffer (200 mM NaH_2_PO_4_ pH 6.8, 5% (w/v) SDS, 1 mM EDTA, 8 M Urea, 1 mg brompheno blue, 100 mM DTT), and incubated at 65°C for 15 minutes. The remaining cell lysates were incubated with pre-equilibrated GFP-Trap agarose beads (Chromotek) for 30-60 minutes at 4°C rotating. Two washes were performed with GFP pulldown buffer, followed by four additional washes in GFP pulldown buffer without detergent on ice. Bound proteins were eluted with 50 μL HU buffer at 65°C for 15 minutes and analyzed by SDS-PAGE and western blotting using an anti-GFP antibody.

### Myo2 GTD-Ldm1 direct binding assay

*E. coli* BL21(DE3 pLysS) cells were transformed with plasmids for co-expression of GST-Ldm1 (pMB363) and GFP-Myo2-GTD(1113-1574) (pMB332) or GST (pMB277) and GFP-Myo2-GTD(1113-1574). Bacteria were pre-cultured overnight in 50 mL LB-medium (2% (w/v) agar, 1% (w/v) Bacto Tryptone, 1% (w/v) Bacto Yeast Extract, 0.5% (w/v) NaCl) containing chloramphenicol, streptomycin and kanamycin (37°C, 280 rpm). In the morning, entire pre-cultures were used to inoculate 1 L of LB-medium containing the same antibiotics and 0.2% glucose. Cells were grown for approximately 6 hours until an OD_600_ of 0.6-0.7 was reached (37°C, 280 rpm). Cultures were cooled on ice to room temperature before induction with 1 mM of isopropyl-beta-D(-)thiogalactopyranoside (IPTG) and further culture at 18°C and 200 rpm for 16 hours. Bacteria were then harvested at 3000 x g for 20 minutes at room temperature. Pellets were washed once in PBS and frozen at -20°C. Prior to lysis cells were thawed on ice and resuspended in 40 mL PBS supplemented with cOmplete protease inhibitor as well as 1 mM phenylmethylsulfonyl fluoride (PMSF). DNAseI and Lysozyme were added and incubated for 30 minutes at 4°C. Lysis was performed by 3 passages of 20000 psi through an LM20 Microfluidizer (Microfluidics). Cell debris was then removed by centrifugation at 25000 x g and 4°C for 30 min. Supernatants (input samples) were incubated with 0.25 mL bed volume equilibrated glutathione Sepharose 4B beads for 3 hours at 4°C. After washing the beads four times with PBS elution was performed in two rounds of 500 µL elution buffer (100 mM Tris-HCl pH 8.0, 10 mM reduced glutathione, 1 mM EDTA pH 8.0). The two corresponding eluate fractions were pooled (eluate samples). Input and eluate samples were analyzed via SDS-PAGE and consecutive immunoblot.

### Proteomic analysis

Proteomic analysis was performed based on Stabile Isotope Labeling by Amino acids in Cell culture (SILAC) ^95^. Briefly, lysine auxotroph yeast cells were inoculated from an overnight preculture and grown in 100 mL SILAC medium to exponential growth phase either in “heavy” lysine ([^13^C_6_^15^N_2_]-lysine) or in “light” lysine. Equal amounts of cells were harvested from both cultures and cell pellets were shock frozen in liquid nitrogen. 500 µL GFP pulldown buffer (150 mM KOAc, 20 mM HEPES pH 7.4, 5 % glycerol, cOmplete protease inhibitor cocktail (Roche) and 1% digitonin) and 500 µL zirconia beads were added to the frozen pellet. Cells were lysed using a FastPrep (MP Biomedicals). Beads were removed by centrifugation and cells were incubated with pre-equilibrated GFP-Trap agarose beads (Chromotek) for 10 minutes at 4°C rotating. Afterwards, beads were washed two times in GFP pulldown buffer and four times in GFP pulldown buffer without detergent at 4°C. Beads from “heavy” and “light” cultures were combined during the last washing step and further processed using the PreOmics sample kit (iST Kit, Preomics) following the “Protocol for iST Sample Preparation Kit for Agarose Immunoprecipitation Samples” except of the digestion with LysC overnight instead of using the provided Trypsin/LysC mixture. The resulting peptides were resuspended in 10 µL LC-Load and 5 µL were subjected to LC-MS/MS analysis.

Reversed-phase chromatography was performed on a Thermo Ultimate 3000 RSLCnano system connected to a Q ExactivePlus mass spectrometer (Thermo) through a nano-electrospray ion source ^96^. Peptides were separated on a 50 cm PepMap C18 easy spray column (Thermo) with a column temperature of 35°C. Peptides were eluted from the column at a constant flow rate of 200 nL/min. Therefore, a linear gradient of acetonitrile starting with 12% buffer B (20% H_2_0, 80% acetonitrile) in 0.1% formic acid which was increased to 35% buffer B for 70 minutes was used, followed by a 20 minute increase to 60% buffer B and finally 10 minutes to reach 90% buffer B. Eluted peptides from the column were directly electro-sprayed into the mass spectrometer. Mass spectra were acquired on the Q ExactivePlus in a data-dependent mode to automatically switch between full scan MS and up to ten data-dependent MS/MS scans. The maximum injection time for full scans was 50 ms, with a target value of 3,000,000 at a resolution of 70,000 at m/z = 200. The ten most intense multiply charged ions (z ≥ 2) from the survey scan were selected with an isolation width of 1.6 Th and fragment with higher energy collision dissociation with normalized collision energies of 27. Target values for MS/MS were set at 100,000 with a maximum injection time of 80 ms at a resolution of 17,500 at m/z = 200. To avoid repetitive sequencing, the dynamic exclusion of sequenced peptides was set at 20 s. The resulting MS and MS/MS spectra were analyzed using MaxQuant (version 2.5.0.0, www.maxquant.org/; ^97,98^). All calculations and plots were performed with the R software package (www.r-project.org/; RRID:SCR_001905). The mass spectrometry proteomics data have been deposited to the ProteomeXchange Consortium via the PRIDE partner repository with the dataset identifier PXD060814 and 10.6019/PXD060814 ^99^.

### Structure prediction and visualization

Structure predictions for the complex of Myo2-GTD and Ldm1 were calculated with AlphaFold 3 ^100^ and conservation was analyzed with the ConSurf Web Server ^101,102^. ChimeraX ^103^ was utilized for superposition of experimental and predicted structures, electrostatic surface potential calculations and figure preparation.

### Quantification and statistical analysis

Fluorescence microscopy images were processed and quantified using ImageJ. Background values of each image were subtracted after measuring the image modal grey value. All visibly budded cells were considered for analysis and categorized according to their bud size, which was defined as the percentage of the daughter cell area relative to the mother cell area. Four bud size categories were considered: I, 0-24%; II, >24-39%; III, >39-50%; IV, >50-61%. For analysis of organelle distribution between daughter and mother cells (Figures 2A-F, 3I, 5F and G, 6C, 6F, 6H, S2C, S2E), the ratio of the mean intensity signal of the respective fluorescent organelle marker in the daughter cell (I_D_) over the mother cells (I_M_) was calculated (I_D_/I_M_). For analysis of secretory vesicles (Figure 2G), I_D_/I_M_ represents the ratio of the mean fluorescent signal of the vesicular structure in the bud to the mean fluorescent signal of the cytoplasm in the mother cell. For Ldm1 enrichment in polarized cell regions (Figure 2I), two circular ROIs were drawn at the polarized site and in the cytosolic area. Subsequently, the ratio of the maximal signal intensity obtained for each ROI was calculated. For quantification of the percentage of LDs in the bud (Figure 3B), fluorescently labeled LDs in mother and daughter cells were counted. For quantification of the percentage of mitochondria in the bud (Figure 7B), the fluorescent mitochondrial marker signal was thresholded and the segmented mitochondrial area in the daughter cell was compared to the total segmented mitochondrial area.

To quantify the effect of Myo2-Fis1 and Ldm1 on mitochondrial distribution in *Δypt11 mmr1-ts* mutants (Figure 7E, S5E), daughter cells containing fluorescent mitochondrial marker signal were counted. To determine the mobile LD fraction (Figure 3D), individual LDs were counted and evaluated manually over time series of 60 s.

All quantifications were performed based on at least three independent experiments. The number of cells analyzed is indicated in each figure legend. In bar graphs, data represents the mean ± standard deviation (SD). In box graph, whiskers represent the 5-95 percentile. T-tests were used for comparison between two samples (Figures 2A-G, 2I, 3B, 7B). One-way ANOVA with Bonferroni was used for comparison of multiple samples (Figures 3D, 3I, 5F and G, 6C, 6F, 6H, 7E, S2C, S2E, S5E). Statistical significance was defined as ∗, p<0.05; ∗∗, p<0.01; ∗∗∗, p<0.001. Graphs were created using GraphPad Prism.

### Yeast strains used in this study

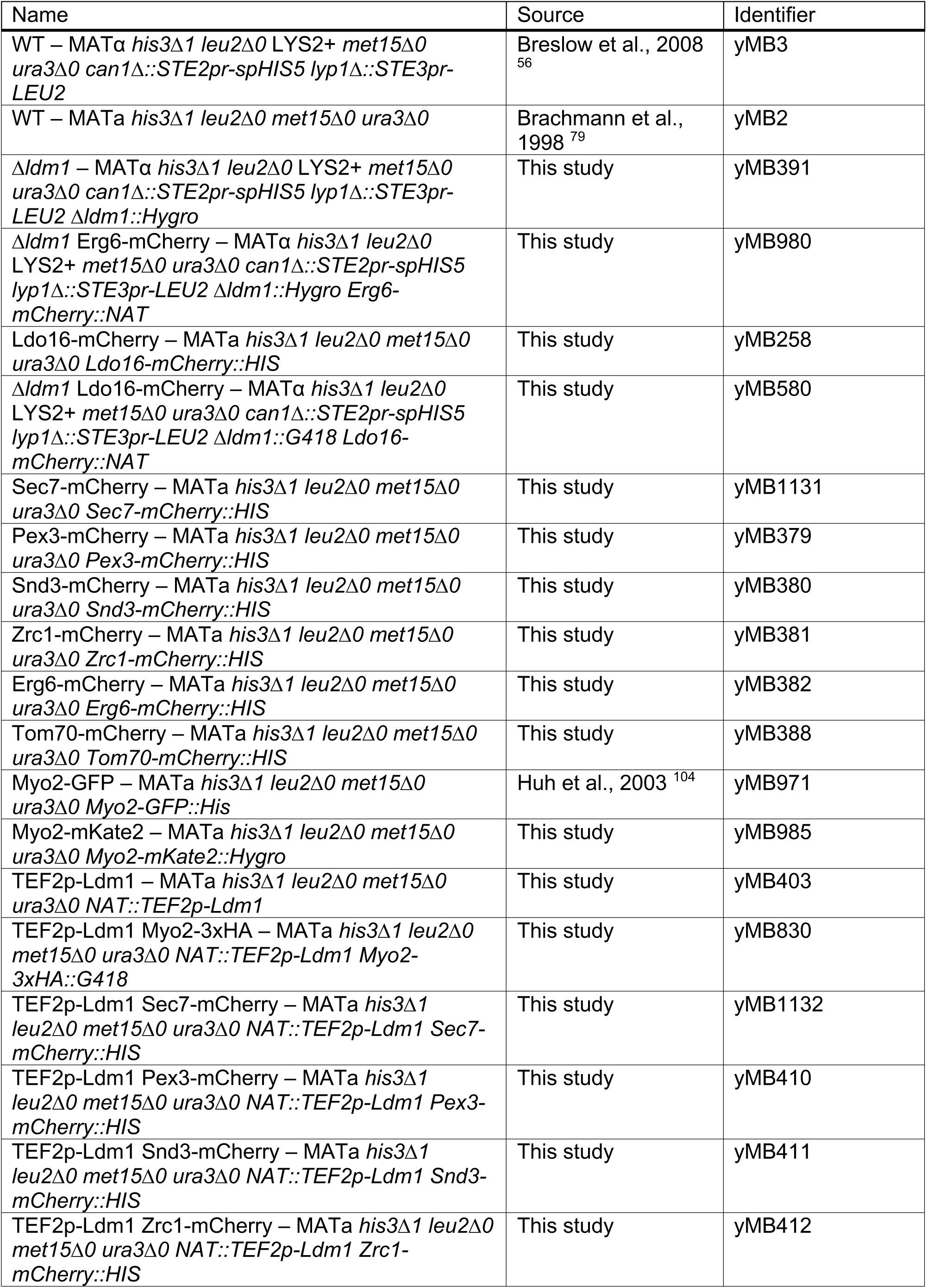
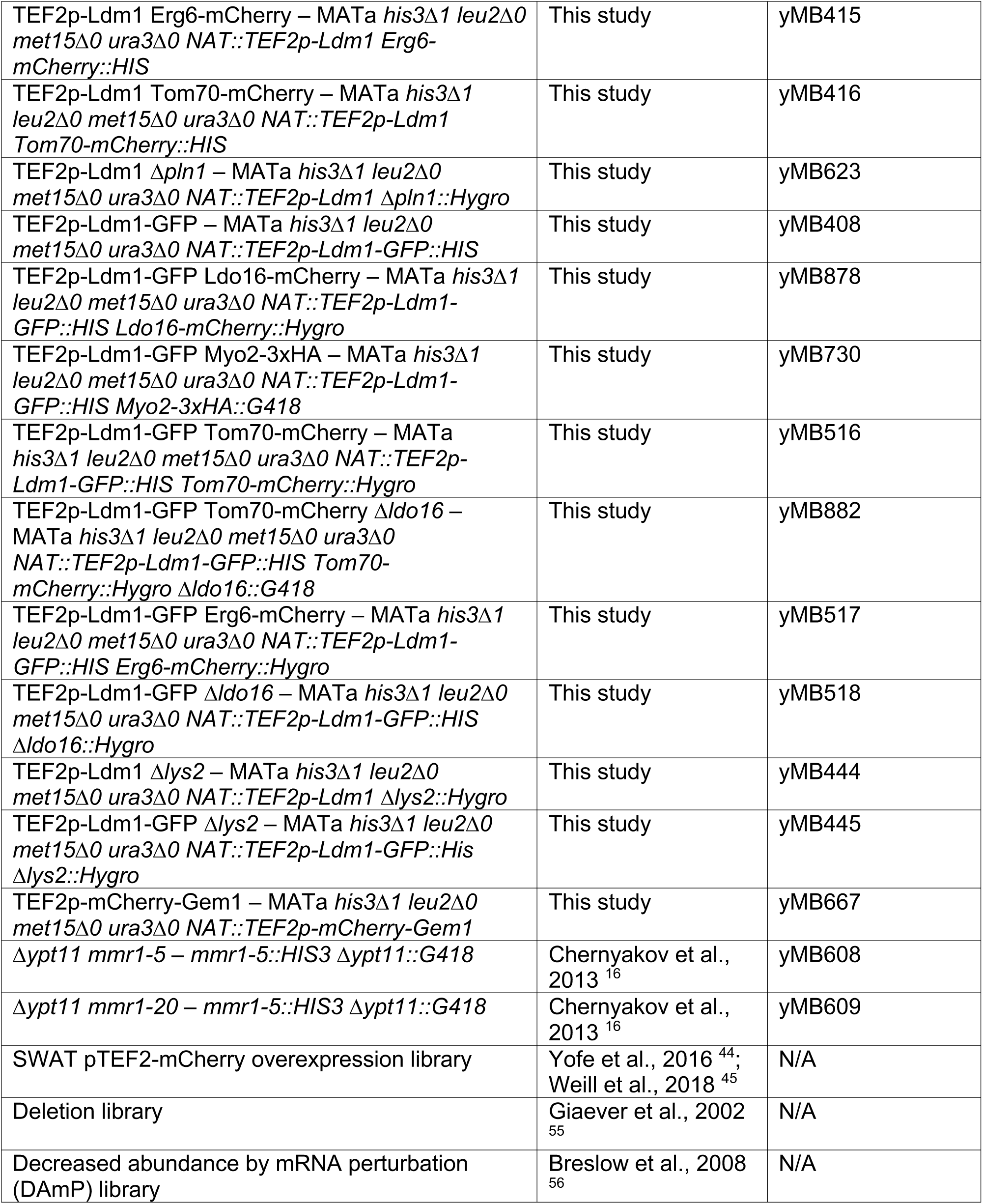

### Plasmids used in this study

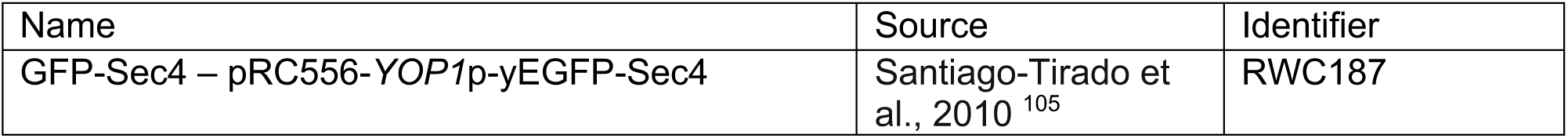
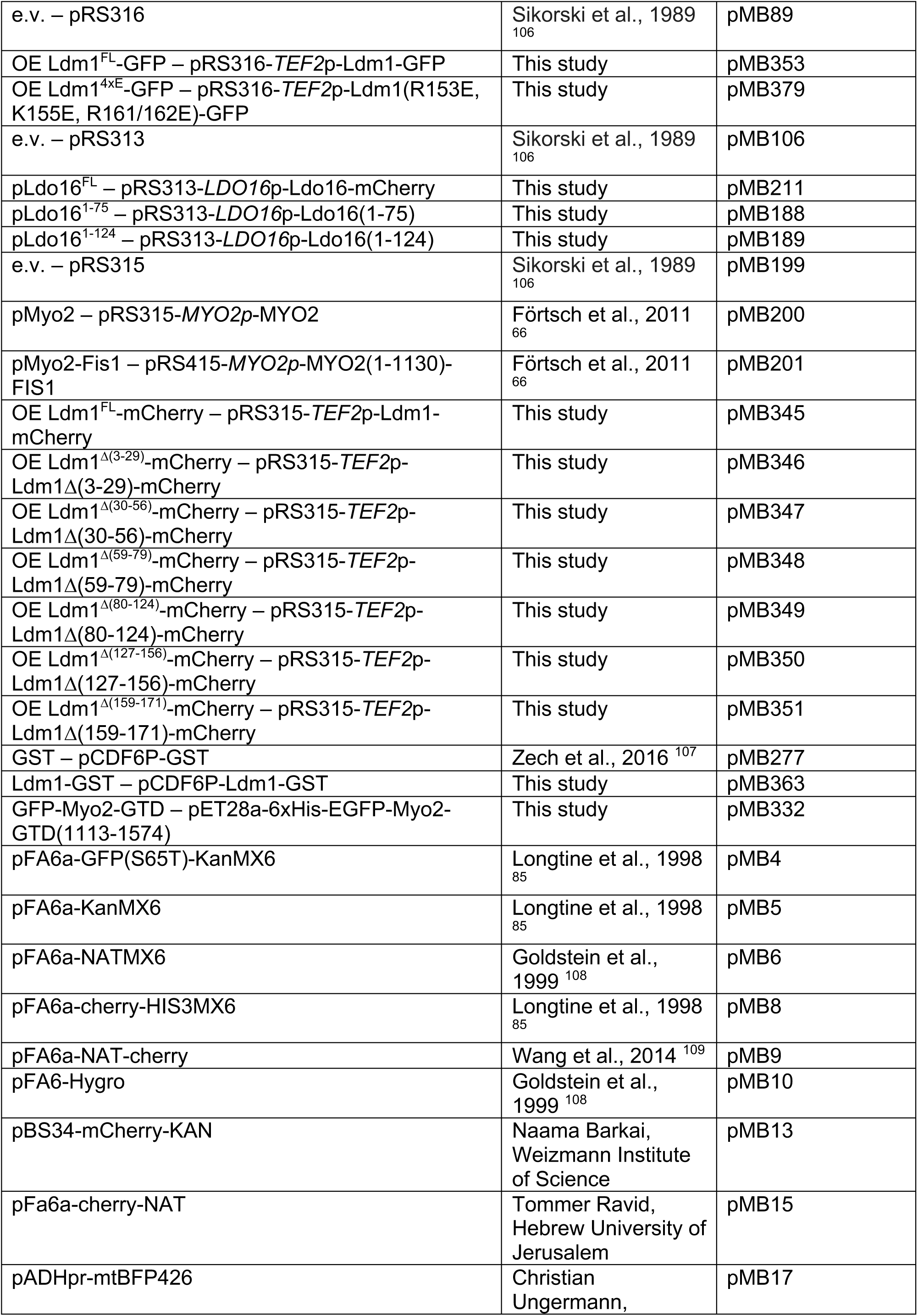
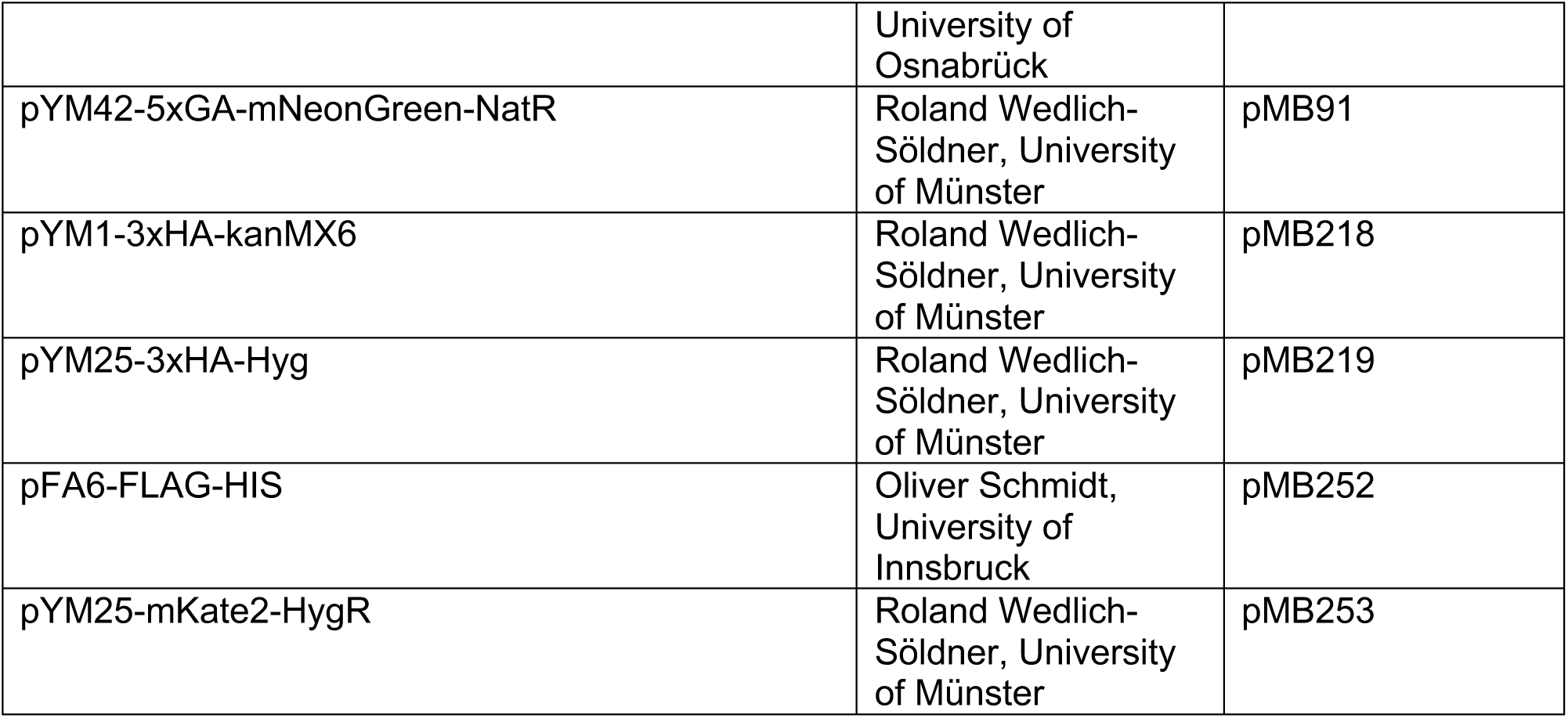

### Primers used in this study

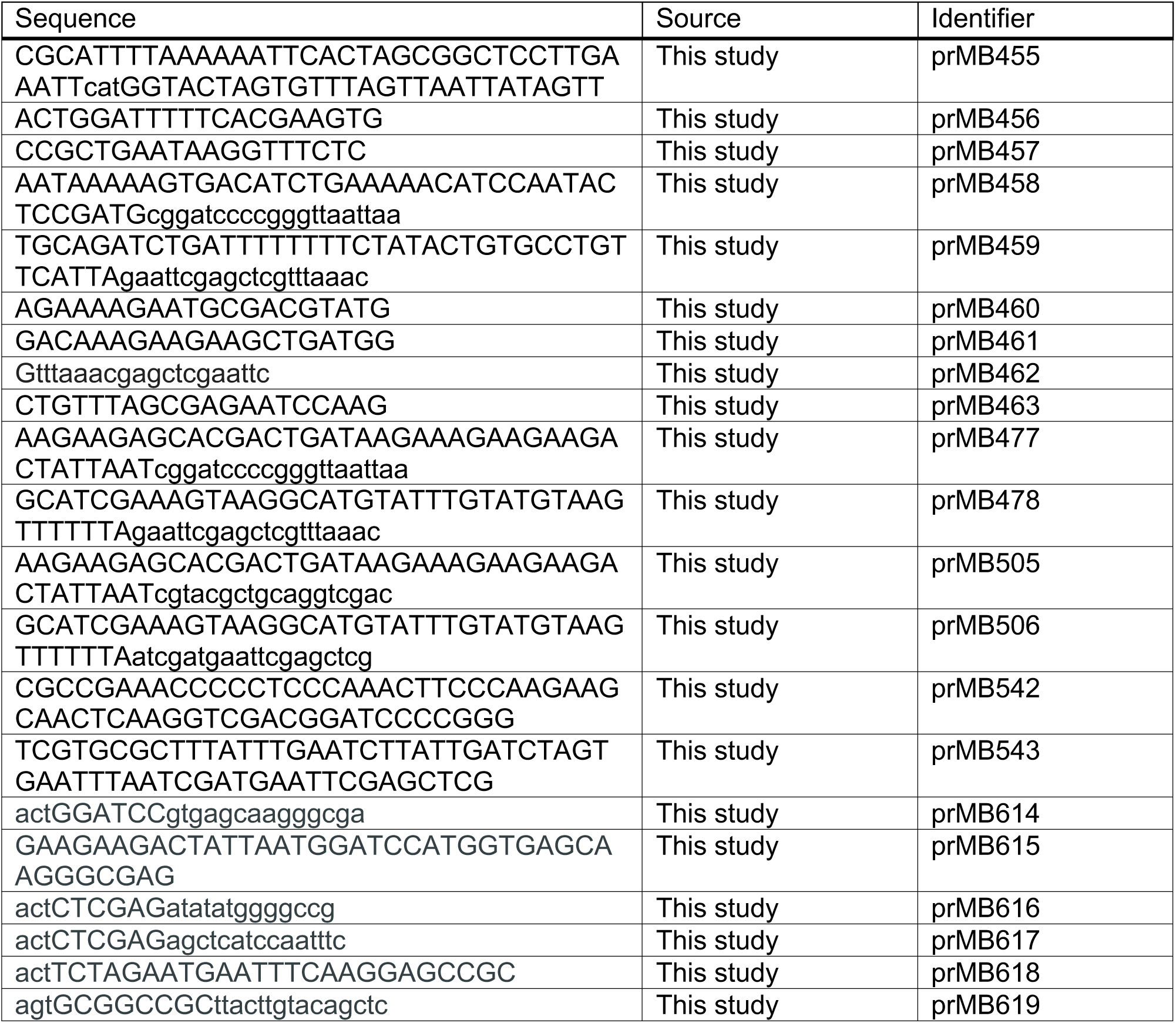
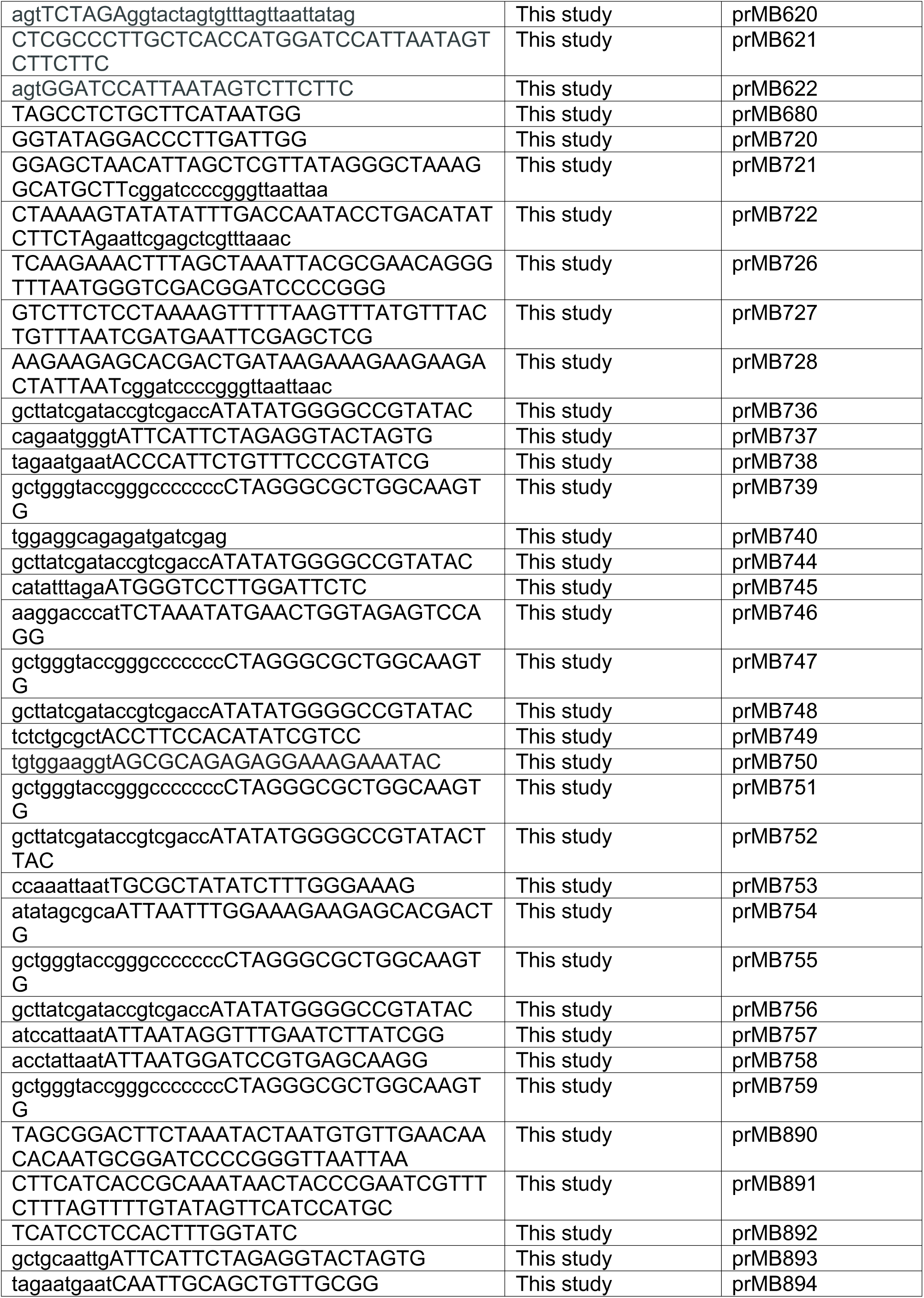
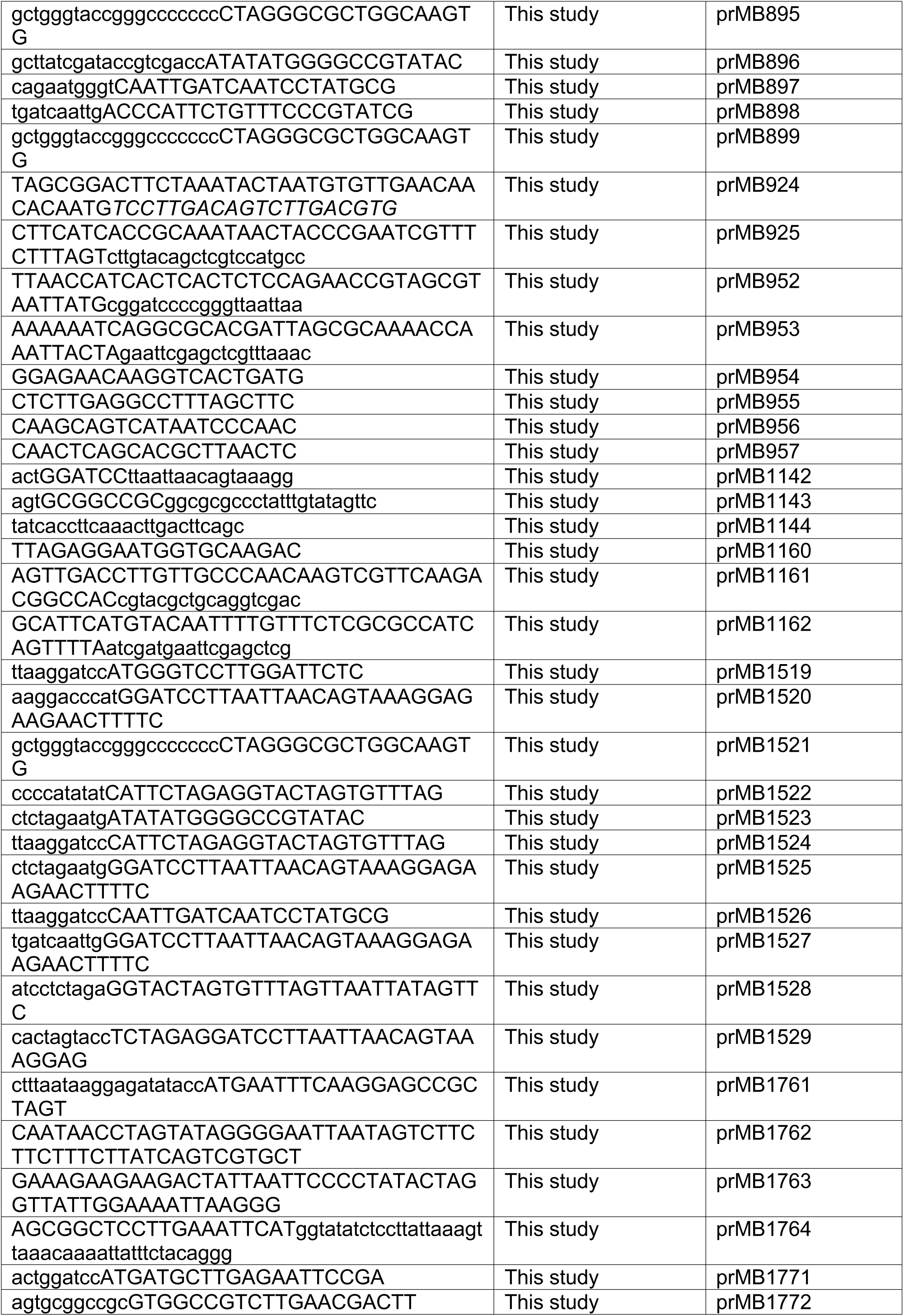
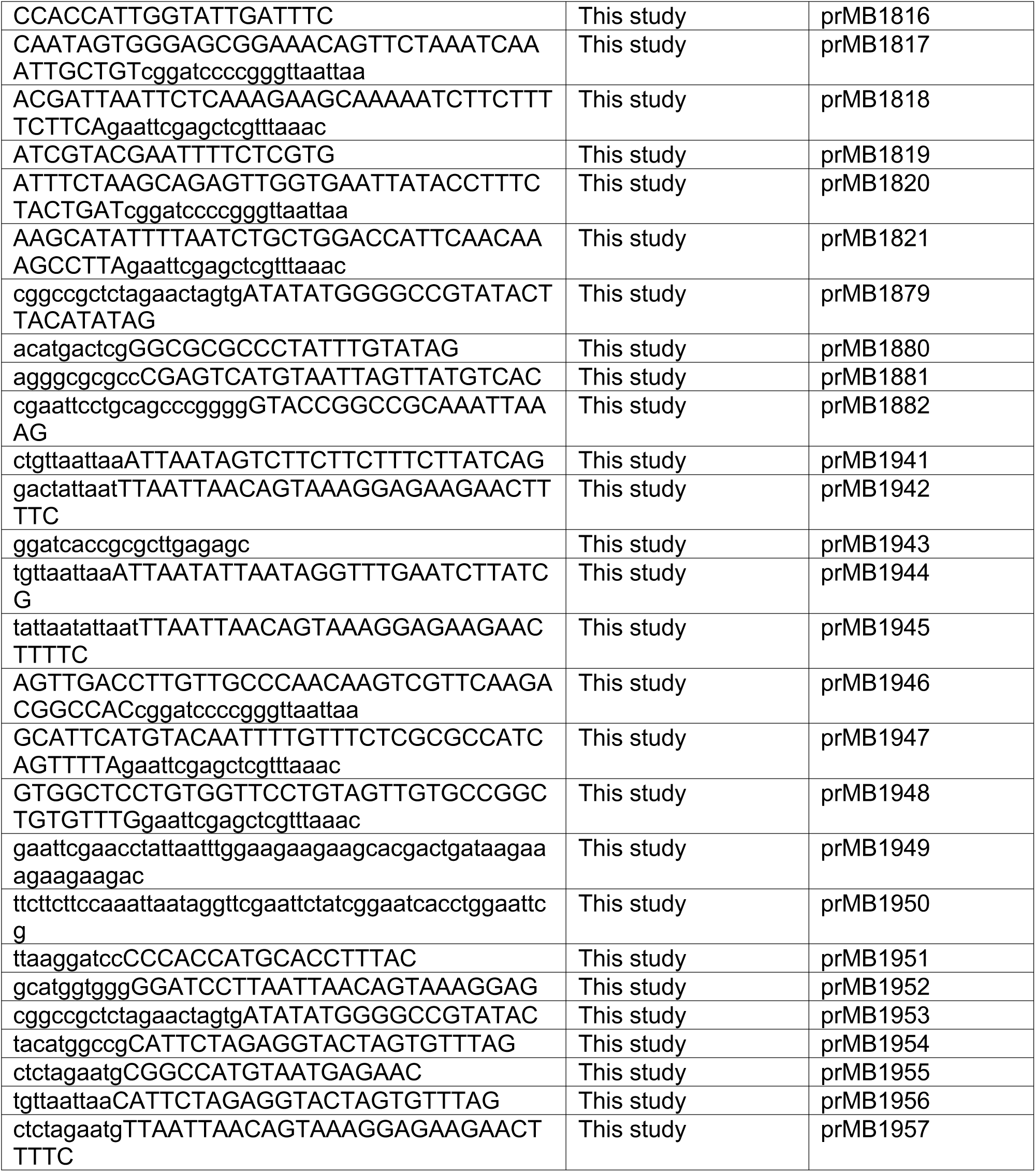

### Antibodies used in this study

| Name | Source | Identifier |
| --- | --- | --- |
| Ldm1 | This study | aMB64 |
| Ldo16 | Diep et al., 2024<sup>41</sup> | aMB13 |
| Om45 | Nikolaus Pfanner,<br>University of<br>Freiburg | 1390-2 |
| Pln1 | Nikolaus Pfanner,<br>University of<br>Freiburg | 1987-7 |
| Tom70 | Nikolaus Pfanner,<br>University of<br>Freiburg | 657-5 |
| Tom20 | Nikolaus Pfanner,<br>University of<br>Freiburg | 3225-7 |
| Tom40 | Nikolaus Pfanner,<br>University of<br>Freiburg | 5104-5 |
| Tim54 | Nikolaus Pfanner,<br>University of<br>Freiburg | 2012-4 |
| GAPDH | Thermo Fischer | Cat # MA5-15738;<br>RRID:<br>AB_10977387 |
| GFP | Roche | Cat#11814460001 |
| HA | Proteintech | Cat# 51064-2-AP;<br>RRID:AB_110423<br>21 |
| GST | Sigma-Aldrich | Cat# G7781;<br>RRID:AB_259965 |
| anti-mouse IgG | Dianova | Cat#115-035-003 |
| anti-rabbit IgG | Dianova | Cat#111-035-003 |
| anti-mouse IgG | Sigma | Cat#A4416;<br>RRID:AB_258167 |
| anti-rabbit IgG | Sigma | Cat#A0545;<br>RRID:AB_257896 |

### Software used in this study

| Name | Source | Identifier |
| --- | --- | --- |
| Image J | Schneider et al.,<br>2012 <sup>110</sup> | https://imagej.nih.gov/ij/download.html<br>RRID:SCR_003070 |
| Adobe Illustrator | Adobe Inc. | RRID:SCR_010279 |
| GraphPad | GraphPad Software Inc. | Prism 5.0<br>RRID:SCR_002798 |
| IMOD 4.11 | Mastronarde et al.,<br>2017 <sup>90</sup> | http://bio3d.colorado.edu/imod<br>RRID:SCR_003297 |
| SerialEM 4.0 | Mastronarde et al.,<br>2005 <sup>88</sup> | http://bio3d.colorado.edu/SerialEM/<br>RRID:SCR_017293 |
| MotionCor2 | Zheng et al., 2017 <sup>89</sup> | https://emcore.ucsf.edu/cryoem-software<br>RRID:SCR_016499 |
| MATLAB | Mathworks | Version 2019b<br>http://www.mathworks.com/products/matlab/<br>RRID:SCR_001622 |
| tom_deconv | Tegunov et al., 2019 <sup>91</sup> | https://github.com/dtegunov/tom_deconv |
| MemBrain-Seg | Lamm et al., 2024 <sup>92</sup> | https://github.com/teamtomo/membrain-seg |
| Chimera | Eric F Pettersen, et al., 2004 <sup>93</sup> | RRID:SCR_004097 |
| MaxQuant (version 2.5.0.0) | Cox and Mann, 2008 <sup>98</sup>; Cox et al., 2011 <sup>97</sup> | www.maxquant.org/ |
| R software package | www.r-project.org/ | RRID:SCR_001905 |
| AlphaFold 3 | Abramson et al., 2024 <sup>100</sup> | RRID:SCR_025885 |
| RSCB-PDB | Berman et al., 2002 <sup>111</sup> | RRID:SCR_012820 |
| Amira | Thermo Fisher Scientific | Version 2022.1<br>https://www.thermofisher.com/de/de/home/electron-microscopy/products/software-em-3d-vis/amira-software.html<br>RRID:SCR_014305 |
| ChimeraX | Meng, et al., 2023 <sup>103</sup> | RRID:SCR_015872 |
| ConSurf | Yariv et al., 2023 <sup>102</sup>, Landau et al., 2005 <sup>101</sup> | https://consurf.tau.ac.il/consurf_index.php |

## Notes

### Competing Interest Statement

The authors have declared no competing interest.

