## Supplement for "The Myo2 adaptor Ldm1 and its receptor Ldo16 mediate actin-dependent lipid droplet motility"

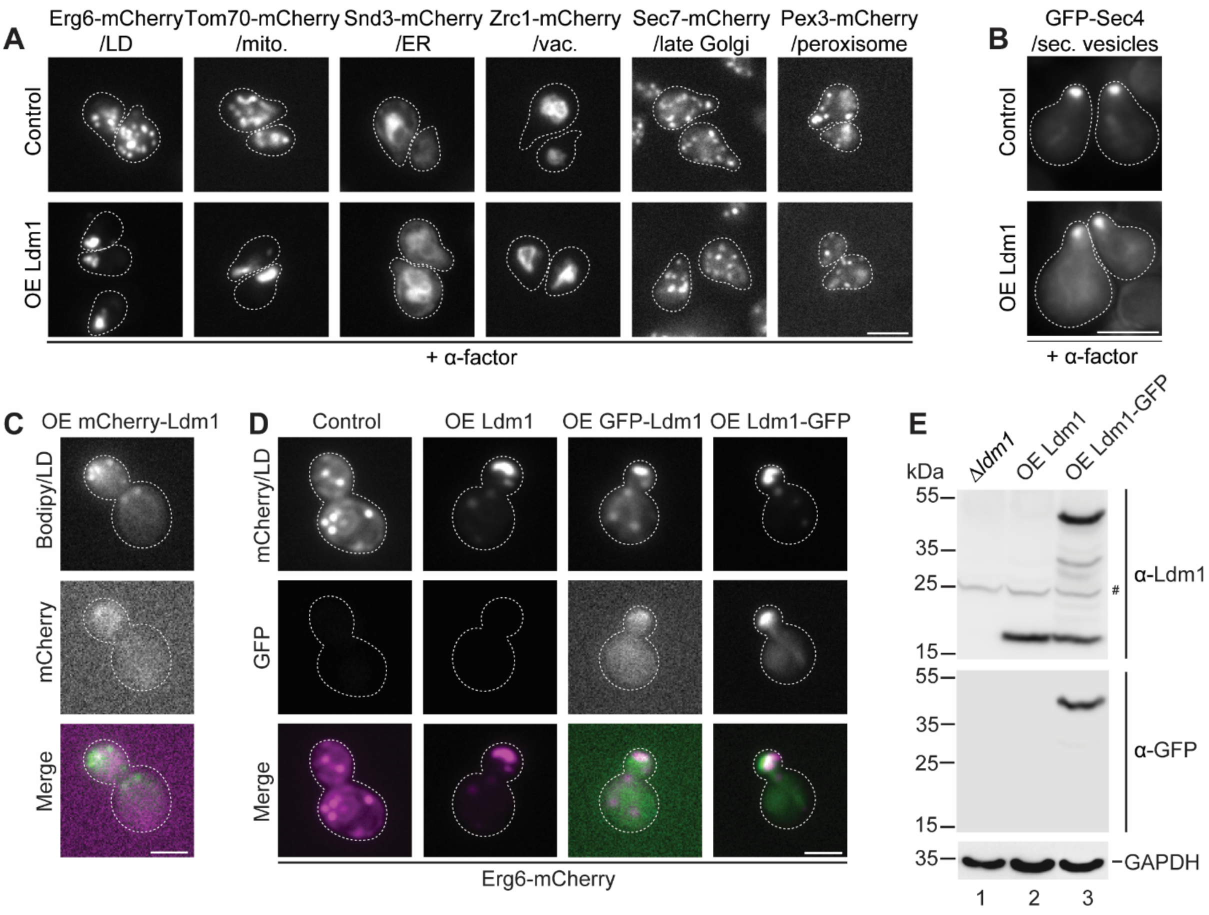


**Figure S1. Characterization of Ldm1 overexpression strains. Related to Figure 2.**

(A) Ldm1 overexpressing (*TEF2* promoter) and control cells expressing mCherry-fused organelle markers were treated for 2 hours with the mating pheromone α-factor and analyzed by microscopy. Scale bar, 5 μm.

(B) Secretory vesicles were labeled by plasmid-derived GFP-Sec4 in control and p*TEF2*-Ldm1 cells and analyzed as in (A).

(C) p*TEF2*-mCherry-Ldm1 cells were stained with BODIPY493/503. Scale bar, 5 μm.

(D) Control, p*TEF2*-Ldm1, p*TEF2*-GFP-Ldm1, p*TEF2*-Ldm1-GFP cells expressing Erg6-mCherry were analyzed by microscopy. Scale bar, 5 μm.

(E) Proteins from *Δldm1*, p*TEF2*-Ldm1 and p*TEF2*-Ldm1-GFP cells were extracted and subjected to SDS-PAGE and western blotting using α-Ldm1, α-GFP and α-GAPDH antibodies. #, unspecific band.

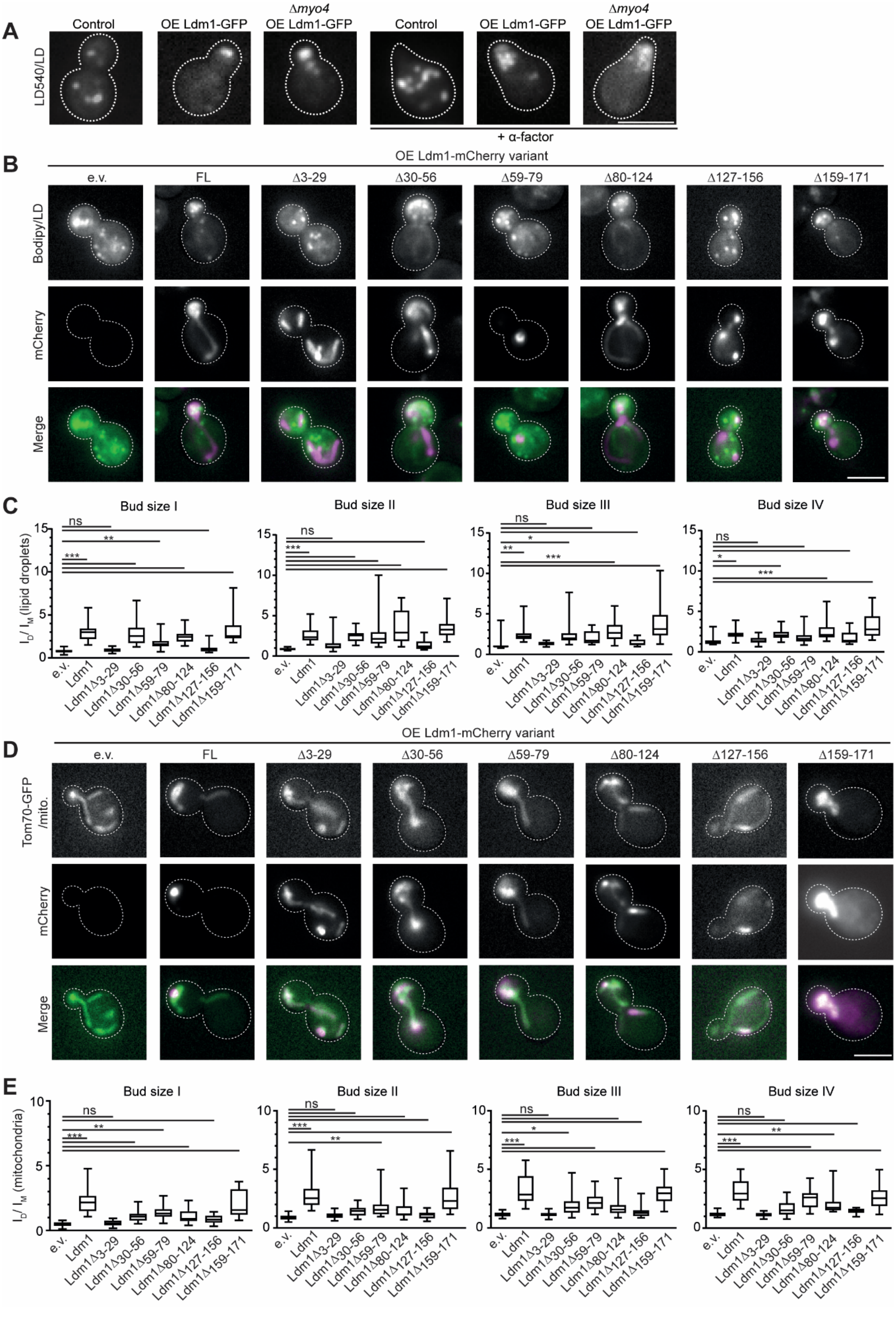

A
Control
OE Ldm1-GFP
Δmyo4 OE Ldm1-GFP
Control
OE Ldm1-GFP
Δmyo4 OE Ldm1-GFP
LD540/LD
+ α-factor
B
OE Ldm1-mCherry variant
e.v.
FL
Δ3-29
Δ30-56
Δ59-79
Δ80-124
Δ127-156
Δ159-171
Bodipy/LD
mCherry
Merge
C
Bud size I
Bud size II
Bud size III
Bud size IV
ID / IM (lipid droplets)
ns
*
**
***
e.v.
Ldm1
Ldm1Δ3-29
Ldm1Δ30-56
Ldm1Δ59-79
Ldm1Δ80-124
Ldm1Δ127-156
Ldm1Δ159-171
D
OE Ldm1-mCherry variant
e.v.
FL
Δ3-29
Δ30-56
Δ59-79
Δ80-124
Δ127-156
Δ159-171
Tom70-GFP /mito.
mCherry
Merge
E
Bud size I
Bud size II
Bud size III
Bud size IV
ID / IM (mitochondria)
ns
*
**
***
e.v.
Ldm1
Ldm1Δ3-29
Ldm1Δ30-56
Ldm1Δ59-79
Ldm1Δ80-124
Ldm1Δ127-156
Ldm1Δ159-171

**Figure S2. Structure function analysis of Ldm1. Related to Figure 5.**
(A) Control, p*TEF2*-Ldm1-GFP and p*TEF2*-Ldm1-GFP *Δmyo4* cells were treated for 2 hours with the mating pheromone $\alpha$-factor and were stained with the red neutral lipid dye LD540. Scale bar, 5 µm.
(B) Control cells transformed with plasmids for expression of indicated Ldm1-mCherry variants (*TEF2* promoter) were stained with BODIPY493/503 and analyzed by microscopy. e.v., empty vector: Scale bar, 5 µm.
(C) Mean intensity ratio of fluorescent LD marker signal in daughter over mother cells ($I_D/I_M$) is depicted as boxplots (5-95 percentile). Cells were categorized by bud size: I, 0-24%, II, >24-39%, III, >39-50%, IV, >50-61%. e.v., empty vector. ns, not significant; $*$, p<0.05; $**$, p<0.01; $***$, p<0.001. Scale bar, 5 µm. N=60 cells, n=3.
(D) Tom70-GFP cells transformed with plasmids for expression of indicated Ldm1-mCherry variants (*TEF2* promoter) were analyzed by microscopy. e.v., empty vector: Scale bar, 5 µm.
(E) Tom70-GFP cells were analyzed as in (C) to assess mitochondrial ($I_D/I_M$). e.v., empty vector. ns, not significant; $*$, p<0.05; $**$, p<0.01; $***$, p<0.001. Scale bar, 5 µm. N=60 cells, n=3.

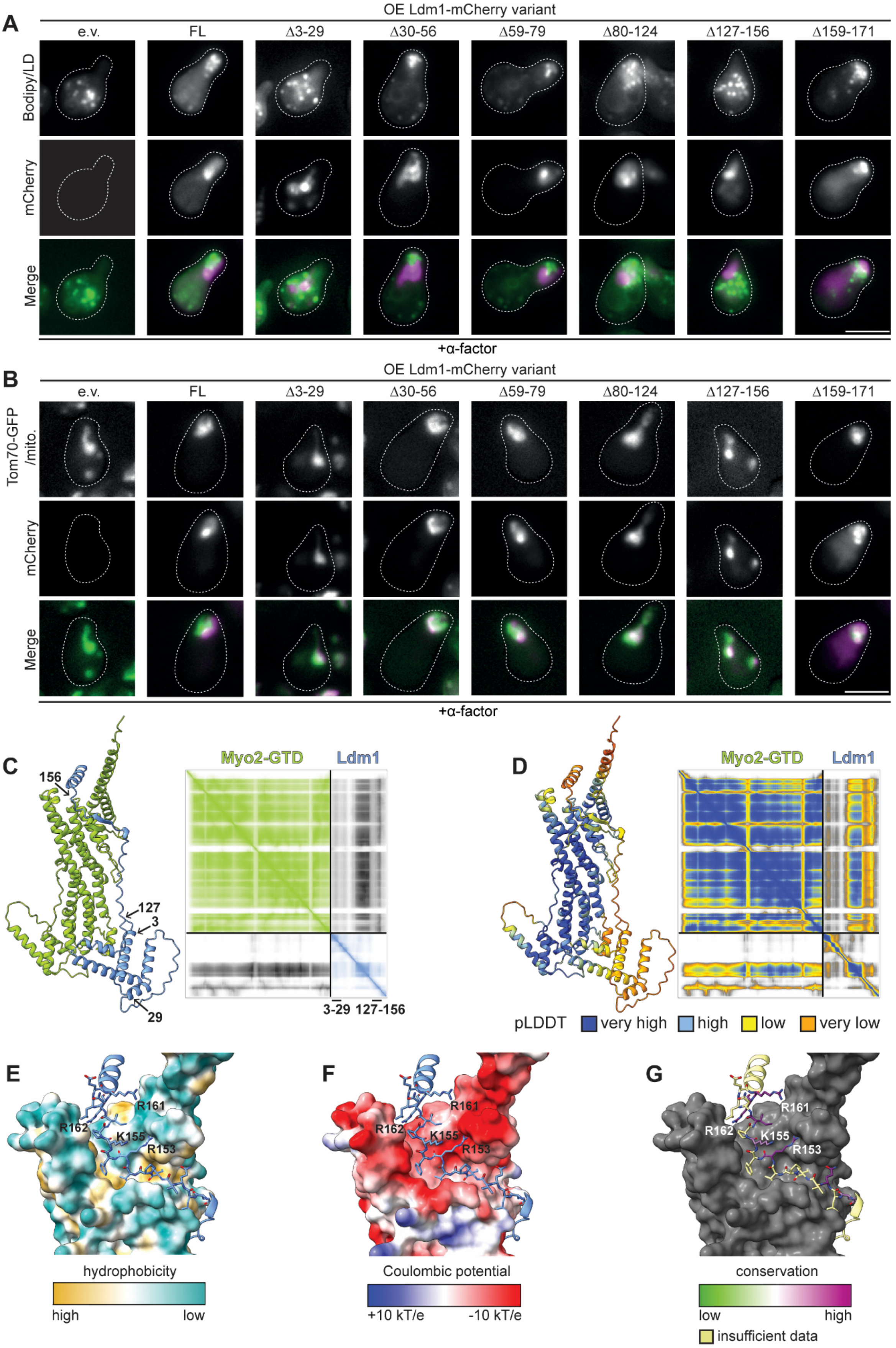
A
OE Ldm1-mCherry variant
e.v.
FL
Δ3-29
Δ30-56
Δ59-79
Δ80-124
Δ127-156
Δ159-171
Bodipy/LD
mCherry
Merge
+α-factor
B
OE Ldm1-mCherry variant
Tom70-GFP /mito.
mCherry
Merge
+α-factor
C
156
127
3
29
Myo2-GTD
Ldm1
3-29
127-156
D
Myo2-GTD
Ldm1
pLDDT
very high
high
low
very low
E
R161
R162
K155
R153
hydrophobicity
high
low
F
R161
R162
K155
R153
Coulombic potential
+10 kT/e
-10 kT/e
G
R161
R162
K155
R153
conservation
low
high
insufficient data

**Figure S3. Interplay between Ldm1 and Myo2. Related to Figure 5.**
(A) Control cells transformed with plasmids for expression of indicated Ldm1-mCherry variants (*TEF2* promoter) were treated for 2 hours with $\alpha$-factor, stained with BODIPY493/503 and analyzed by microscopy. e.v., empty vector. Scale bar, 5 µm.
(B) Tom70-GFP cells transformed with plasmids for expression of indicated Ldm1-mCherry variants (*TEF2* promoter) were treated for 2 hours with $\alpha$-factor and were analyzed by microscopy. e.v., empty vector. Scale bar, 5 µm.
(C) AlphaFold 3 structure prediction of Myo2-GTD (green) in complex with Ldm1 (blue) with corresponding PAE (predicted aligned error) plot colored by chain (ptm = 0.75, iptm = 0.71). Domain boundaries chosen for interaction site mapping are labelled.
(D) Same as (C), but colored by predicted local distance difference test (pLDDT) scores.
(E-G) Close ups of interface A between Myo2-GTD (surface representation) and Ldm1 (cartoon). The adaptor protein binds Myo2-GTD at a hydrophobic cleft framed by negatively charged residues, engaging with hydrophobic and highly conserved arginine (R153, R161, R162) and lysine residues (K155).

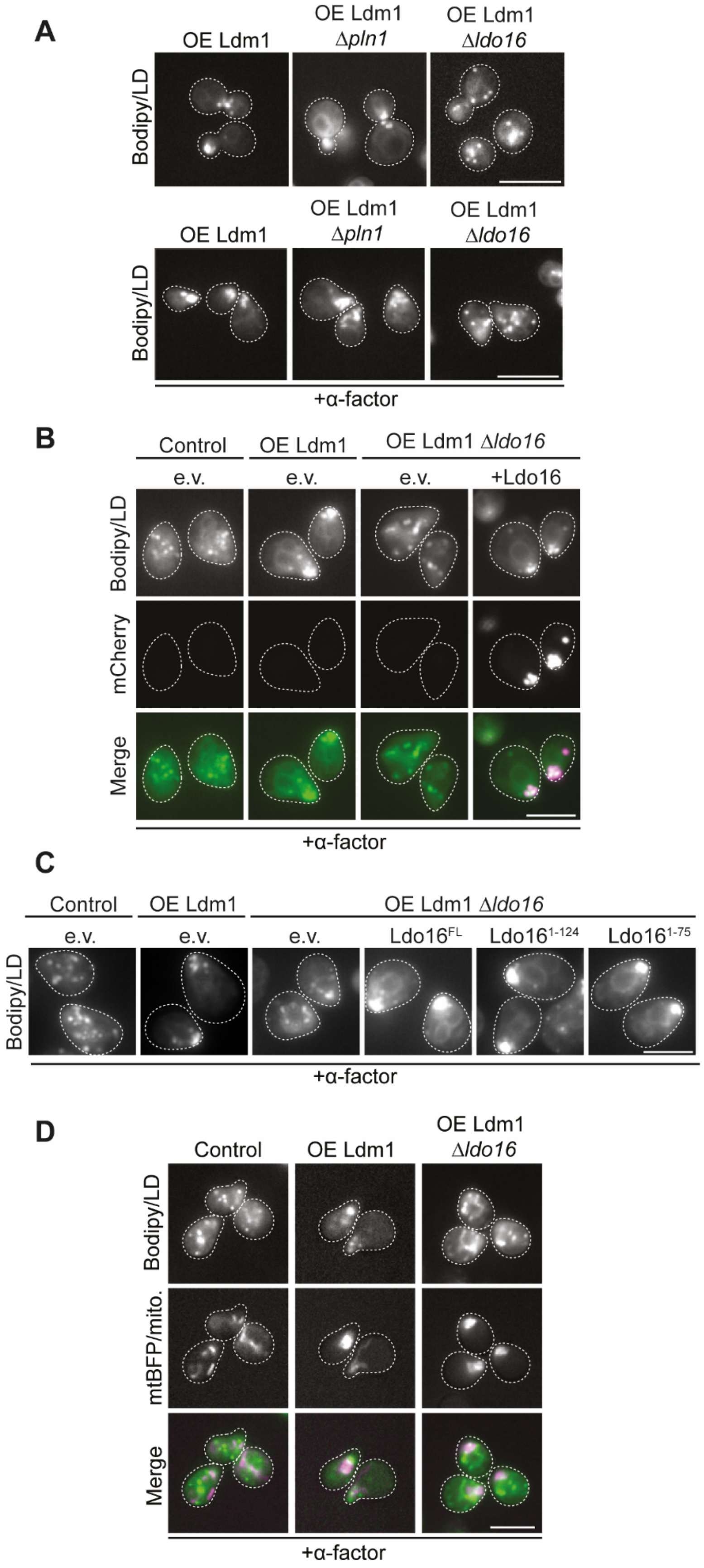

A
OE Ldm1
OE Ldm1 Δpln1
OE Ldm1 Δldo16
Bodipy/LD
OE Ldm1
OE Ldm1 Δpln1
OE Ldm1 Δldo16
Bodipy/LD
+α-factor
B
Control
OE Ldm1
OE Ldm1 Δldo16
e.v.
e.v.
e.v.
+Ldo16
Bodipy/LD
mCherry
Merge
+α-factor
C
Control
OE Ldm1
OE Ldm1 Δldo16
e.v.
e.v.
e.v.
Ldo16FL
Ldo161-124
Ldo161-75
Bodipy/LD
+α-factor
D
Control
OE Ldm1
OE Ldm1 Δldo16
Bodipy/LD
mtBFP/mito.
Merge
+α-factor

**Figure S4. Ldo16 is an Ldm1 partner protein. Related to Figure 6.**

(A) p*TEF2*-Ldm1-GFP, p*TEF2*-Ldm1-GFP *Δpln1* and p*TEF2*-Ldm1-GFP *Δldo16* cells were treated for 2 hours with α-factor or left untreated, stained with BODIPY493/503 and analyzed by microscopy. Scale bar, 5 µm.

(B) Control, p*TEF2*-Ldm1, and p*TEF2*-Ldm1 *Δldo16* cells were transformed with a centromeric plasmid for expression of Ldo16 from its own promoter (+Ldo16) or an empty vector (e.v.) and treated for 2 hours with α-factor and stained with BODIPY493/503. Scale bar, 5 µm.

(C) Control, p*TEF2*-Ldm1, and p*TEF2*-Ldm1 *Δldo16* cells were transformed with indicated plasmids, treated for 2 hours with α-factor and stained with BODIPY493/503. FL, centromeric plasmid for expression of full length Ldo16-mCherry from the p*LDO16* promoter. 1-124 and 1-75, centromeric plasmids for expression of truncated Ldo16 variants comprising amino acids 1-124 or 1-75 from the p*LDO16* promoter. e.v., empty vector. Scale bar, 5 µm.

(D) Control, p*TEF2*-Ldm1, and p*TEF2*-Ldm1 *Δldo16* cells were transformed with a plasmid for expression of mtBFP to label mitochondria, treated for 2 hours with α-factor and stained with BODIPY493/503. Scale bar, 5 µm.

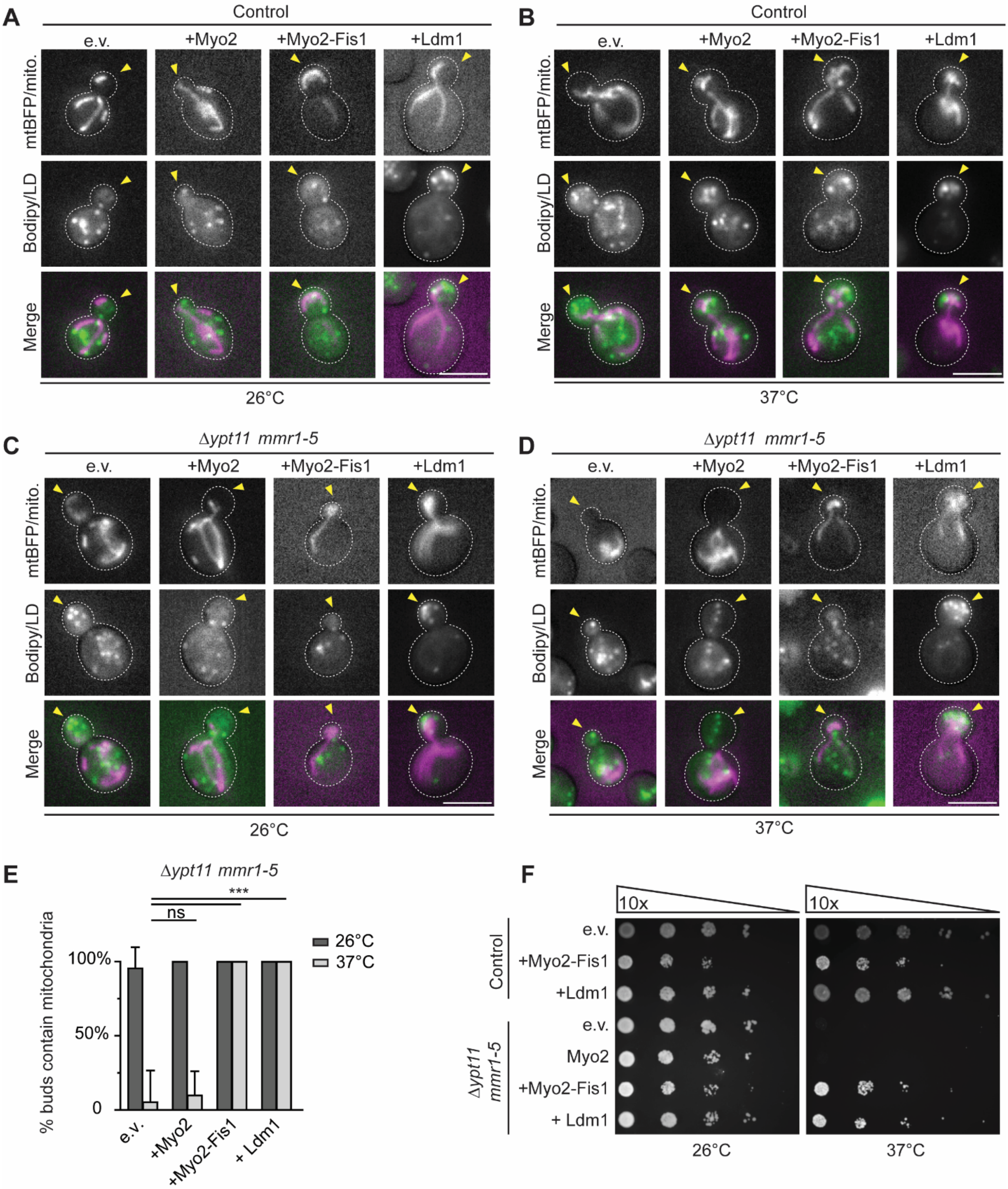

A
Control
e.v.
+Myo2
+Myo2-Fis1
+Ldm1
mtBFP/mito.
Bodipy/LD
Merge
26°C
B
Control
e.v.
+Myo2
+Myo2-Fis1
+Ldm1
mtBFP/mito.
Bodipy/LD
Merge
37°C
C
Δypt11 mmr1-5
e.v.
+Myo2
+Myo2-Fis1
+Ldm1
mtBFP/mito.
Bodipy/LD
Merge
26°C
D
Δypt11 mmr1-5
e.v.
+Myo2
+Myo2-Fis1
+Ldm1
mtBFP/mito.
Bodipy/LD
Merge
37°C
E
Δypt11 mmr1-5
***
ns
% buds contain mitochondria
100%
50%
0
26°C
37°C
e.v.
+Myo2
+Myo2-Fis1
+ Ldm1
F
10x
10x
Control
e.v.
+Myo2-Fis1
+Ldm1
Δypt11 mmr1-5
e.v.
Myo2
+Myo2-Fis1
+ Ldm1
26°C
37°C

**Figure S5. Ldm1 function is independent of Ypt11 and Mmr1. Related to Figure 7.**
(A) Control cells expressing mtBFP for labeling of mitochondria were transformed with plasmids for overexpression of Myo2 (+Myo2), a Myo2 variant fused to the outer mitochondrial membrane protein Fis1 (+Myo2-Fis1) for attachment of the motor protein to mitochondria independent of Myo2-adaptor proteins, or Ldm1 (+Ldm1). Cells were cultured at 26°C, stained with BODIPY493/503, and analyzed by microscopy. e.v., empty vector. Scale bar, 5 µm.
(B) Same as in (A), with the difference that cells were incubated at 37°C for 1.5 hours prior to and during imaging.
(C) *Δypt11 mmr1-5* cells expressing mtBFP for labeling of mitochondria were transformed with plasmids described in (A). Cells were cultured at 26°C, stained with BODIPY493/503, and analyzed by microscopy. e.v., empty vector. Scale bar, 5 µm.
(D) Same as in (C), with the difference that cells were incubated at 37°C for 1.5 hours prior to and during imaging.
(E) The percentage of dividing cells with mitochondria present in the bud was determined. Data represented as mean ± SD. ns, not significant; ∗∗∗, $p<0.001$. $N\geq60$ cells, n=3.
(F) Control and *Δypt11 mmr1-5* cells transformed with plasmids described in (A) were grown overnight at 26°C, adjusted to an $OD_{600}$ of 0.05, serially 10-fold diluted, spotted on agar plates and grown at 26°C or 37°C for 3 days. e.v., empty vector.

| Phenotype | Name | ORF | Description |
|---|---|---|---|
| Polarized LDs | *LDM1* | *YER085C* | Putative protein of unknown function |
| | *RCY1* | *YJL204C* | F-box protein involved in recycling endocytosed proteins |
| Less LDs | *BCK2* | *YER167W* | Serine-threonine-rich protein involved in PKC1 signaling |
| | *DGK1* | *YOR311C* | Diacylglycerol kinase localized to the ER |
| | *IZH4* | *YOL101C* | Membrane protein involved in zinc ion homeostasis |
| | *SET4* | *YJL105W* | Chromatin-associated protein regulating stress response |
| | *SFP1* | *YLR403W* | Regulator of ribosomal protein gene expression |
| | *UGX2* | *YDL169C* | Protein of unknown function |
| More LDs | *AEP1* | *YMR064W* | Protein required for expression of F1F0 ATPase subunit 9 |
| | *BOI1* | *YBL085W* | Protein involved in polar growth |
| | *CSE4* | *YKL049C* | Centromeric histone H3-like protein |
| | *HPT1* | *YDR399W* | Dimeric hypoxanthine-guanine phosphoribosyltransferase |
| | *NGR1* | *YBR212W* | RNA binding protein that negatively regulates growth rate |
| | *NUP1* | *YOR098C* | FG-nucleoporin component of the nuclear pore complex |
| | *PIF1* | *YML061C* | DNA helicase |
| | *RAD27* | *YKL113C* | 5' to 3' exonuclease, 5' flap endonuclease |
| | *SET1* | *YHR119W* | Histone methyltransferase, subunit of the COMPASS |
| | *SPO11* | *YHL022C* | Meiosis-specific protein that initiates meiotic recombination |
| | *TIM11* | *YDR322C-A* | Subunit e of mitochondrial F1F0-ATPase |
| | *VIK1* | *YPL253C* | Subunit of a kinesin-14 heterodimeric motor with Kar3 |
| | *YDL129W* | *YDL129W* | Protein of unknown function |
| | *YOR019W* | *YOR019W* | Protein of unknown function |
| | *YOR072W-B* | *YOR072W-B* | Putative protein of unknown function |
| | *ZDS1* | *YMR273C* | Regulator of Swe1-dependent polarized growth |
| Strong clustering | *CSF1* | *YLR087C* | Protein with similarity to lipid transfer protein Vps13 |
| | *FMP27* | *YLR454W* | Protein with similarity to lipid transfer protein Vps13 |
| | *LDO16* | *YMR148W* | Seipin partner, LD-vacuole tether together with Vac8 |
| | *MTC4* | *YBR255W* | Protein of unknown function |
| | *PUL4* | *YNR063W* | Putative zinc-cluster protein |
| Partial clustering | *ACE2* | *YLR131C* | Transcription factor required for septum destruction |
| | *FAA1* | *YOR317W* | Long chain fatty acyl-CoA synthetase |
| | *ICY1* | *YMR195W* | Protein of unknown function |
| | *MEX67* | *YPL169C* | Poly(A)RNA binding protein for nuclear mRNA export |
| | *NIP100* | *YPL174C* | Large subunit of the dynactin complex |
| | *OLE1* | *YGL055W* | Delta(9) fatty acid desaturase |
| | *RGD1* | *YBR260C* | RhoGAP for Rho3 and Rho4 |
| | *RTC4* | *YNL254C* | Protein of unknown function |
| | *SCT1* | *YBL011W* | Acyltransferase for glycerolipid synthesis |
| | *SLD2* | *YKL108W* | Replication initiation protein |
| | *SPO77* | *YLR341W* | Protein required for spore wall formation |
| | *SPT23* | *YKL020C* | Regulator of *OLE1* transcription |
| | *STP4* | *YDL048C* | Predicted transcription factor |
| | *TLD1* | *YDR275W* | Protein that regulates lipolysis |

**Table S1. List of genes overexpression of which results in altered abundance, morphology and distribution of LDs. Related to Figure 1.**

| Phenotype | Name | ORF | Description |
|---|---|---|---|
| Loss of LD polarization | *ARF1* | *YDL192W* | GTPase of the Ras superfamily |
| | *AVO2* | *YMR068W* | TORC2 subunit |
| | *BNI1* | *YNL271C* | Formin that nucleates formation of linear actin filaments |
| | *BRE5* | *YNR051C* | Ubiquitin protease cofactor |
| | *CAF16* | *YFL028C* | Part of CCR4-NOT regulatory complex |
| | *CDC50* | *YCR094W* | Endosomal partner of phospholipid flippase Drs2p |
| | *CHO2* | *YGR157W* | Phosphatidylethanolamine methyltransferase (PEMT) |
| | *CIK1* | *YMR198W* | Kinesin-associated protein |
| | *DRS2* | *YAL026C* | Trans-Golgi network phospholipid flippase |
| | *FAT1* | *YBR041W* | Very long chain fatty acyl-CoA synthetase |
| | *FOX2* | *YKR009C* | Multifunctional enzyme of peroxisomal beta-oxidation |
| | *INO4* | *YOL108C* | Transcription factor involved in phospholipid synthesis |
| | *KCS1* | *YDR017C* | IP6 and IP7 kinase |
| | *LDB18* | *YLL049W* | Component of the dynactin complex |
| | *LDO16* | *YMR148W* | Seipin partner, LD-vacuole tether together with Vac8 |
| | *MRPS12* | *YNR036C* | Mitochondrial ribosomal protein |
| | *NCA3* | *YJL116C* | Regulator of expression of F1F0 ATPase subunits 6 and 8 |
| | *OPI9* | *YLR338W* | Dubious open reading frame, overlaps with *VRP1* |
| | *PUB1* | *YNL016W* | Poly (A)+ RNA-binding protein |
| | *RAD51* | *YER095W* | Strand exchange protein involved in DNA DSB breaks |
| | *RAD52* | *YML032C* | Protein involved in homologous recombination |
| | *RPS27B* | *YHR021C* | Protein component of the small ribosomal subunit |
| | *SEC26* | *YDR238C* | Essential beta-coat protein of the COPI coatomer |
| | *TEC1* | *YBR083W* | Transcription factor |
| | *VAM3* | *YOR106W* | Syntaxin-like vacuolar t-SNARE |
| | *VMA4* | *YOR332W* | V-ATPase subunit E |
| | *VPS4* | *YPR173C* | AAA-ATPase involved in MVB protein sorting |
| | *VPS52* | *YDR484W* | Component of the GARP complex |
| | *VPS53* | *YJL029C* | Component of the GARP complex |
| | *VPS61* | *YDR136C* | Dubious open reading frame, overlaps with *RGP1* |
| | *VPS63* | *YLR261C* | Putative protein of unknown function, overlaps with *YPT6* |
| | *YKR073C* | *YKR073C* | Putative protein of unknown function |
| | *YPT6* | *YLR262C* | Rab family GTPase |
| | *YPT7* | *YML001W* | Rab family GTPase |

**Table S2. List of genes deletion of which results in loss of Ldm1-dependent LD accumulation in polarized cell regions. Related to Figure 4.**
